# Climate predicts *w*Mel *Wolbachia* frequency variation in *Drosophila melanogaster*, but genomic variation reflects a recent incomplete cytoplasmic sweep

**DOI:** 10.64898/2026.05.22.727337

**Authors:** Nitin Ravikanthachari, Emily L. Behrman, Jack Beltz, William R. Conner, Paul R. Schmidt, Brandon S Cooper

## Abstract

Maternally transmitted *Wolbachia* occupy roughly half of terrestrial arthropod species, but the factors maintaining their variable population frequencies are poorly understood. In *Drosophila melanogaster*, the *w*Mel *Wolbachia* variant occurs at intermediate and variable frequencies globally. We document rapid *w*Mel frequency shifts up to 0.33 between consecutive weeks in experimental and natural orchards in the eastern United States, with frequencies peaking at intermediate temperatures and declining at thermal extremes. Seven years of seasonal sampling at a Pennsylvania orchard showed *w*Mel frequencies consistently higher in summer than fall, consistent with temperature-dependent maternal transmission. Bayesian models applied to 248 locations across five continents and 42 years of sampling corroborated the previously described *w*Mel frequency cline in eastern Australia but found no latitudinal pattern on any other continent. Precipitation seasonality and driest-quarter precipitation were the strongest global predictors, absorbing the among-continent *w*Mel frequency variation that latitude left unexplained. Wet-season temperature predicted *w*Mel frequency in Australia, where the wettest quarter coincides with summer and peak host reproduction. Analysis of 339 individually sequenced *w*Mel genomes identified 38 *w*Mel SNPs associated with latitude, but these associations did not persist after accounting for cytoplasmic lineage structure, and 35 of the 38 were private to a remnant southwestern European lineage. Our results establish that local climatic conditions shape *w*Mel frequencies globally, plausibly through effects on maternal transmission fidelity that depend on the seasonal alignment of warm temperatures with host reproduction. *w*Mel genomic variation, in contrast, reflects the incomplete replacement of ancestral *w*MelCS by *w*Mel, rather than local adaptation.

## Introduction

*Wolbachia* endosymbionts occur in a large fraction of arthropod species (Werren and Windsor 2000; Hilgenboecker et al. 2008; Zug and Hammerstein 2012; Weinert et al. 2015) and vary widely in frequency within and among host populations (Hoffmann 1988; Hamm et al. 2014; Ahmed et al. 2015; Kriesner et al. 2016; Cooper et al. 2017; Wheeler et al. 2021; Turelli et al. 2022; Lečić et al. 2024). This variation arises because the parameters governing *Wolbachia* frequency dynamics – maternal transmission fidelity, net fitness effects on hosts, and the strength of *Wolbachia*-induced reproductive effects (Hoffmann et al. 1990) – may each be sensitive to environmental and genetic context (Kriesner et al. 2016; Shropshire et al. 2020; Hague et al. 2022; Hoffmann and Cooper 2024). Cytoplasmic incompatibility (CI), in which *Wolbachia*-modified sperm kill embryos lacking the endosymbiont, is the most common *Wolbachia* reproductive effect, occurring in roughly half of all *Wolbachia* infections (Yen and Barr 1973; Shropshire et al. 2020; Turelli et al. 2022). CI varies widely in strength (Shropshire et al. 2020; Shropshire et al. 2022), which influences equilibrium *Wolbachia* frequencies in host populations and their propensity to vary. For example, *w*Ri causes strong CI (Hoffmann et al. 1986), contributing to its rapid spread to high and relatively stable frequencies in worldwide *D. simulans* populations (Kriesner et al. 2013). In contrast, *w*Mel causes weak CI in *D. melanogaster* unless males are very young (Reynolds and Hoffmann 2002; Shropshire et al. 2021), and persists at lower and more variable frequencies than *w*Ri (Hoffmann et al. 1998; Kriesner et al. 2016). We assess *w*Mel frequency variation at multiple spatial and temporal scales, and in the context of its incomplete replacement of the ancestral *w*MelCS variant (Riegler et al. 2005).

Kriesner et al. (2016) provided the most comprehensive analysis of *w*Mel frequency variation to date, using more than two decades of sampling to document a persistent *w*Mel frequency cline in eastern Australia. Applying the equilibrium model of Hoffmann et al. (1990), they proposed that fitness costs during reproductive dormancy in colder temperate regions could reduce the net fitness advantage of carrying *w*Mel and plausibly explain lower observed *w*Mel frequencies at higher Australian latitudes. Hague et al. (2022) subsequently showed that cool rearing temperatures reduced *w*Mel abundance at the posterior pole of developing oocytes and the fidelity of maternal transmission. Mathematical modeling indicated these transmission differences, combined with weak CI and a conjectured fitness benefit of ∼25%, could generate the frequency differences observed between tropical and temperate Australian populations. Analysis of natural genotypes – both tropical and temperate – and reciprocal introgressions indicated contributions of host and *w*Mel genomes to the temperature dependence of *w*Mel transmission fidelity (Hague et al. 2022).

However, the cline observed in eastern Australia did not generalize to other continents—no latitudinal pattern was evident in Eurasia, and the cline was shallower in eastern North America, where Kriesner et al. (2016) proposed that seasonal recolonization from lower latitudes or local refugia may decouple observed frequencies from transmission-selection equilibria. Tropical frequencies were relatively high among sparsely sampled African populations, but frequencies were well below the expected equilibrium in some equatorial populations. Even within the interannually stable Australian cline (Kriesner et al. 2016), monthly sampling of a Gold Coast population had previously revealed marked frequency fluctuations, including a drop from ∼0.92 to ∼0.25 across a single monthly interval (Hoffmann et al. 1998). Whether *w*Mel frequencies in *D. melanogaster* follow distinct trajectories at seasonal or shorter timescales, and whether specific environmental factors can explain the among-population frequency variation that latitude cannot, are open questions. That both host and *w*Mel genomes contribute to the temperature dependence of maternal-transmission fidelity implies that *w*Mel genomic variation itself may be structured by environment (Hague et al. 2022).

Richardson et al. (2012) reconstructed complete *w*Mel and mitochondrial genomes from 290 strains and found congruent genealogies organized into six cytoplasmic clades, consistent with strict maternal co-transmission and a single ancestral infection. The most recent common ancestor of all *w*Mel and mitochondrial lineages dates to approximately 8,000 years ago, reflecting a cytoplasmic sweep in which derived *w*Mel lineages have incompletely replaced the ancestral *w*MelCS variant (Riegler et al. 2005; Nunes et al. 2008; Richardson et al. 2012). These clades are geographically structured (Richardson et al. 2012): clade I predominates in North America, clades II and IV in Africa, clade III in Europe and Africa, clade V in Eurasia (Richardson et al. 2012; Ilinsky 2013; Bykov et al. 2019), and a rare clade VIII in Asia (Chrostek et al. 2013; Bykov et al. 2019). Richardson et al. defined clade V from two uninfected French strains based solely on mitochondrial haplotype. Versace et al. (2014) subsequently assembled the first clade V *w*Mel genome from an individually sequenced Portuguese fly. Because *w*Mel and mitochondria are coinherited through the maternal cytoplasm (Richardson et al. 2012), the geographic distribution of *w*Mel clades tracks maternal lineages and the colonization history of *D. melanogaster* (Pool et al. 2012; Sprengelmeyer et al. 2020; Coughlan et al. 2022). Whether *w*Mel genomic variation is associated with environmental conditions independent of the demographic history that shaped its clade structure is uncertain.

Here, we assess environmental predictors of *w*Mel frequency variation at multiple spatiotemporal scales and test whether *w*Mel allele frequencies are shaped by local climate or instead reflect the lineage structure left by the incomplete cytoplasmic sweep. Weekly and seasonal sampling of *D. melanogaster* orchard populations documented rapid frequency shifts associated with temperature, while Bayesian models applied to 248 locations across five continents identified precipitation and temperature predictors that explain the continent-level frequency structure latitude does not capture. We identified 38 *w*Mel SNPs whose derived allele frequencies covaried with latitude, but corrections for lineage structure traced these associations to an early-diverging southwestern European lineage. Apparent environment–allele associations, including those previously interpreted as evidence for temperature-driven selection on individual *w*Mel loci (Hague et al. 2022), reflect this lineage structure. Together, our results establish that local ecological conditions shape *w*Mel frequencies globally, while *w*Mel genomic variation retains the signature of cytoplasmic lineage history. These findings have implications for predicting *Wolbachia* dynamics under environmental change and for *w*Mel-based vector and pest control (Ross, Turelli, et al. 2019; Hoffmann and Cooper 2025).

## Results

### *w*Mel frequency variation in experimental and natural orchard populations

#### Rapid frequency variation in an experimental orchard

We first asked whether *w*Mel frequencies vary on even shorter timescales than previously observed (Hoffmann et al. 1998). We established 12 replicate cage populations in an experimental orchard at the University of Pennsylvania (UPenn) from a genetically diverse outbred stock, sampling replicate cage populations (*N* = 12) weekly from July 7 to October 14, 2023 (*N* = 14 weeks). *w*Mel frequencies did not differ significantly among cages (**Tables S1–S3**) at week one (Firth’s penalized binomial GLM, LRT: χ² = 17.47, df = 11, *P* = 0.095), but they varied significantly across the 14 weeks of sampling (**Tables S1 and S4**; **Fig. S1**; χ² = 37.44, *P* < 0.001). While binomial confidence intervals sometimes overlapped, indicating bouts of temporal stability (*e.g.,* weeks 1–3 and 6–9, **Table S1**), we observed significant frequency changes and sometimes rapid shifts in mean *w*Mel frequency (*e.g.,* week 3: 0.72, 95% CI: 0.64–0.8; week 4: 0.87, 95% CI: 0.81–0.92). *w*Mel frequencies also varied significantly among cages through time (**Fig. S2; Table S4**; χ² = 67.96, *P* < 0.001), with cage-specific frequency trajectories that we examine in the context of temperature below.

#### Rapid and seasonal frequency variation in natural orchards

*w*Mel frequencies also varied significantly through time at both Linvilla Orchard in Media, PA (**Tables S6 and S7**; **Fig. 1A**; χ^2^ = 18.51, *P* = 0.001) and Lohr’s Orchard in Churchville, MD (**Tables S6 and S7**; **Fig. 1A**; χ^2^ = 9.10, *P* = 0.007). At Linvilla, mean frequency initially increased from our first sample in week 2 (0.52, 95% CI: 0.37-0.68) to a maximum frequency of 0.96 (95% CI: 0.85–0.99) in week 5, followed by a rapid decrease to 0.63 (95% CI: 0.48–0.77) in week six that did not differ from our final estimate in week 7 (**Table S6**). This reflected a non-linear frequency trajectory that was supported by a GAM-estimated temporal trend (**Table S7**, **Fig. 1A**, estimated degrees of freedom, EDF = 4.16). *w*Mel frequencies at Lohr’s were relatively stable within each of two sampling bouts (weeks 2–7 and 11–14; binomial GLM slopes nonsignificant, both *P* > 0.8; **Table S5**), with higher frequencies in the second bout. A GAM-estimated temporal trend across all sampled weeks was near-linear (EDF = 1.57), though the data are best interpreted as a step change between two stable periods separated by the sampling gap.

**Figure 1.**
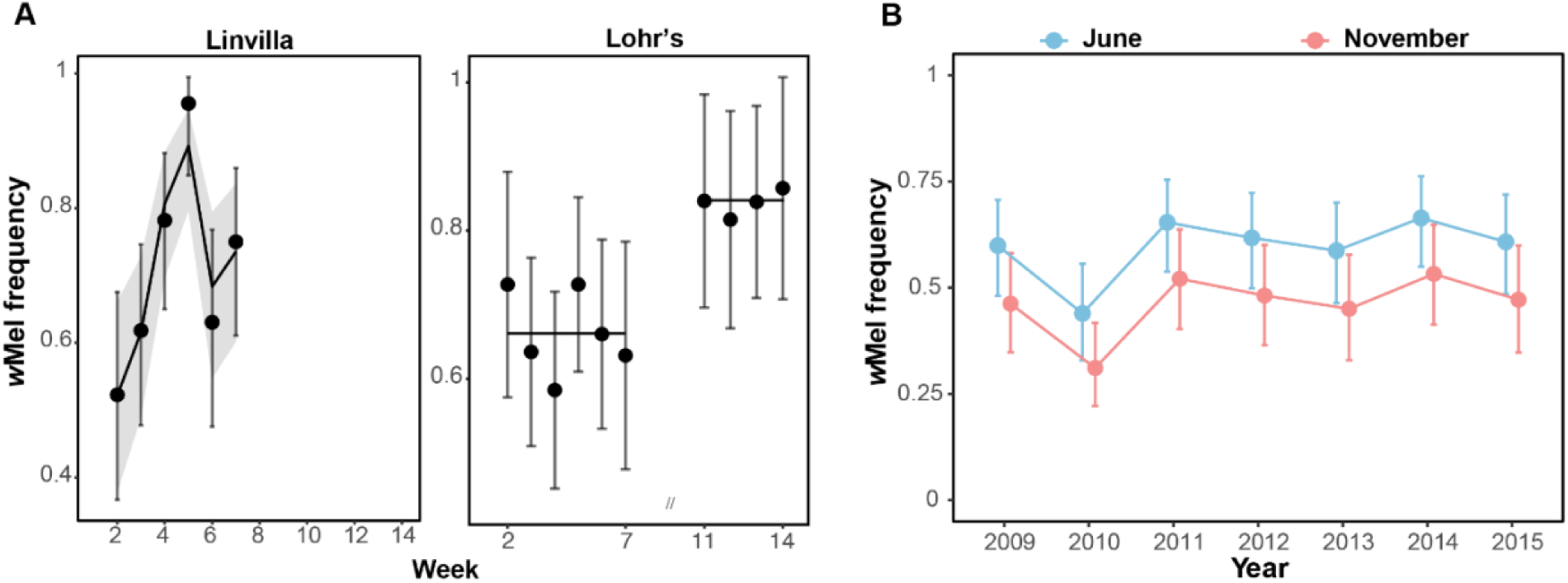
*w*Mel frequencies varied on weekly and seasonal timescales in natural orchards. **(A)** Observed and predicted *w*Mel frequency for the natural orchards at Linvilla, PA and Lohr’s, MD. *w*Mel frequency varied significantly through time at both locations (Linvilla, *P* = 0.001; Lohr’s, *P* = 0.007), but temporal trajectories differed. Points show observed *w*Mel frequencies at each weekly collection with 95% binomial confidence intervals. For Linvilla, the curve and shaded band show the GAM-estimated temporal trend with 95% confidence interval. For Lohr’s, horizontal lines indicate mean *w*Mel frequency within each of two sampling bouts (weeks 2–7 and 11–14), separated by a three-week gap. **(B)** Observed *w*Mel frequencies in June (blue) and November (red) at Linvilla orchard across seven years of seasonal sampling (2009–2015). *w*Mel frequencies were on average ∼0.14 higher in June than November (binomial GLM, est. = −0.55, z = −3.14, *P* = 0.002). This seasonal difference was consistent across years, with no significant month × year interaction (likelihood ratio test, χ² = 8.41, df = 6, *P* = 0.21). Points show observed frequencies for each collection; bars represent 95% binomial confidence intervals.

Seasonal sampling of Linvilla from 2009 to 2015 showed mean *w*Mel frequencies that were, on average, ∼0.14 higher in June than in November (**Table S8**, **Fig. 1B**; est.= −0.55, z value = −3.14, *P =* 0.002). Frequencies were relatively stable across the seven-year period (binomial GLM LRT: χ^2^ = 11.57, *P* = 0.072). Together, our results from the two natural orchards indicate that *w*Mel frequencies varied seasonally and sometimes week-to-week, with temporal trajectories differing between orchards across periods of weekly sampling.

#### Temperature correlates with w*Mel* frequency variation

*w*Mel frequencies were associated with temperature at all three orchards, with frequencies peaking at intermediate temperatures and declining at both thermal extremes (**Fig. 2A–B**). Temperature metrics were lagged by one week, matching the weekly sampling resolution and the mean egg-to-eclosion development time (9.9 days; see Methods). The unimodal signal was robust at lags of 7–10 days but degraded at longer lags, consistent with the temperature–frequency association reflecting conditions during maternal transmission rather than offspring development. At the experimental orchard, *w*Mel frequency varied non-linearly with lagged-weekly mean temperature (LWM; **Table S9A**, quasi-binomial GAM, EDF = 2.60, *P* < 0.001) and lagged-weekly average maximum temperature (LWAM; **Table S9B**, quasi-binomial GAM, EDF = 2.63, *P* < 0.001), but did not differ significantly among cages for either metric (LWM: *P* = 0.57; LWAM: *P* = 0.63). Frequencies peaked at 22.1°C for LWM (peak range: 19.1–25.3°C; **Fig. S3A**) and at 27.3°C for LWAM (peak range: 23.1–30.8°C; **Fig. S3B**).

**Figure 2.**
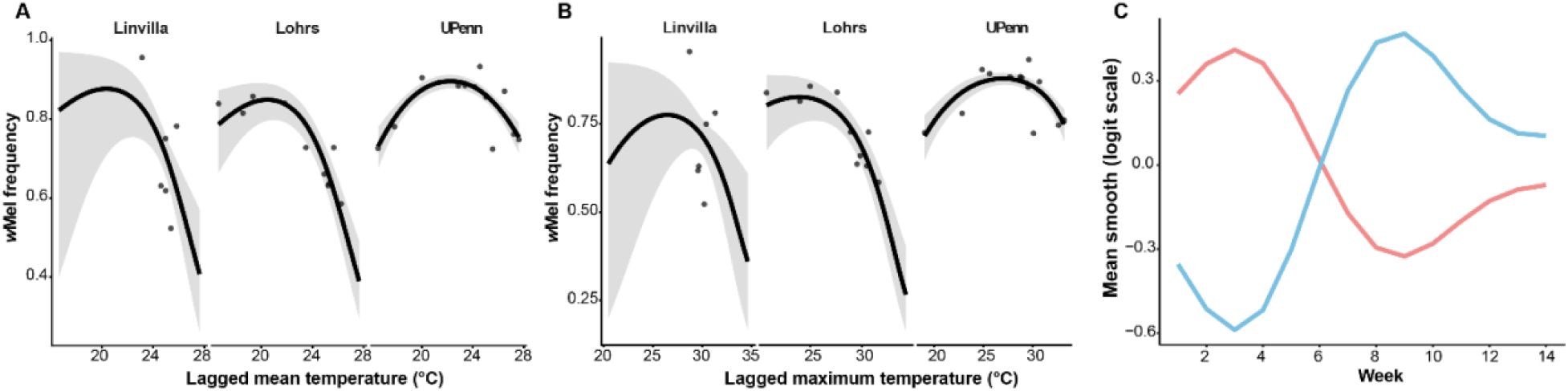
*w*Mel frequency peaks at intermediate temperatures across three orchard populations, with microhabitat shade differences driving opposing trajectories among experimental cages. **(A)** Predicted *w*Mel frequency as a function of lagged weekly mean temperature across all three orchards. Points represent observed weekly *w*Mel frequencies at each location; curves show predicted values from a binomial GAM with location as a fixed effect and a location x temperature interaction smooth allowing the temperature-frequency relationship to differ among sites; temperature values are lagged by one week. *w*Mel frequency was significantly associated with weekly mean temperature (*P* < 0.001), peaking near 20–22°C and declining at both thermal extremes. The strength of this association differed significantly among locations (*P* < 0.001). **(B)** Same model structure as (A) but with lagged weekly average maximum temperature as the predictor. *w*Mel frequency peaked near 24–27°C, with a significant temperature association (*P* < 0.001) and significant among-location differences in the strength of that association (*P* < 0.001). **(C)** Mean estimated temporal trends (logit scale) of *w*Mel frequency over 14 weeks for experimental cages grouped by a priori shade classification: unshaded cages (red; E1–E5, E10, E12) received direct sunlight, while shaded cages (blue; E6–E9, E11) were covered by denser canopy and an adjacent trellis. Shaded and unshaded groups followed opposing frequency trajectories, consistent with the two groups experiencing the same temperature fluctuations at different intensities or timing. Cage-specific temporal trajectories differed significantly (cage × time interaction *P* < 0.001). Trends estimated from a binomial generalized additive model.

The twelve experimental cages differed in sun exposure: cages E1–E5, E10, and E12 were unshaded with direct morning and afternoon sunlight, whereas cages E6–E9 and E11 were shaded by denser canopy and an adjacent trellis. Cages grouped by shade status showed contrasting temporal trajectories (**Fig. 2C**), with within-group correlations substantially higher (mean *r* = 0.49) than between-group correlations (mean *r* = −0.60). In unshaded cages, *w*Mel frequency increased initially until week 2 before declining until week 8, then increasing again from week 9 to week 14. Shaded cages exhibited a mirrored pattern, with frequencies initially decreasing, followed by a rapid increase and subsequent decrease at the same time points. These opposing dynamics are consistent with the two groups experiencing the same temperature fluctuations at different intensities or timing due to microclimate differences.

Across all three orchards, both temperature metrics were significantly associated with *w*Mel frequency (LWM: **Table S10**, binomial GAM, χ² = 59.88, *P* < 0.001; LWAM: **Table S11**, binomial GAM, χ² = 51.65, *P* < 0.001), and the strength of the temperature-frequency association differed among locations for both LWM (**Table S10**, location x temperature interaction: χ² = 17.48, *P* < 0.001) and LWAM (**Table S11**, location x temperature interaction: χ² = 16.26, *P* < 0.001). Peak LWM temperatures were similar at Linvilla (20.3°C; range: 17.1–23.0°C) and Lohr’s (20.5°C; range: 17.4–23.1°C), as were peak LWAM temperatures (Linvilla: 26.4°C, range: 23.3–29.4°C; Lohr’s: 24.0°C, range: 24.0–27.6°C; **Fig. 2A–B**). Baseline *w*Mel frequencies were significantly higher at the experimental orchard than at the natural orchards for LWAM (**Table S11**, UPenn vs. Linvilla: b = 0.84, *P* < 0.001) but not for LWM (**Table S10**, b = 0.51, *P* = 0.05), while the two natural orchards did not differ from each other (Linvilla and Lohr’s: both ∼0.71; LWM *P* = 0.39; LWAM *P* = 0.9). Higher baseline frequencies in the experimental cages may reflect stronger effective CI in enclosed populations where females encounter a higher proportion of young males (Reynolds and Hoffmann 2002; Shropshire et al. 2021). Linvilla estimates spanned large confidence intervals due to sparse observations at lower temperatures and should be interpreted with caution (**Table S12**).

### Global *w*Mel frequency variation

#### Latitude does not explain wMel frequency outside of Australia

Our dataset combined the data compiled by Kriesner et al. (2016), which we refer to as Kdata, with genomic and experimental data collected since their study, yielding frequency estimates from 13,199 individuals across 434 spatiotemporal collections spanning 248 locations on 5 continents and 42 years of sampling (Edata; **Fig. 3A**). We fully replicated Kriesner et al.’s GLM results on the Kdata (**see Supplemental Materials**), but DHARMa diagnostics confirmed significant overdispersion and quantile deviations for all GLMs (**Table S14**). We therefore fit Bayesian beta-binomial models with random intercepts for location and year within location.

**Figure 3.**
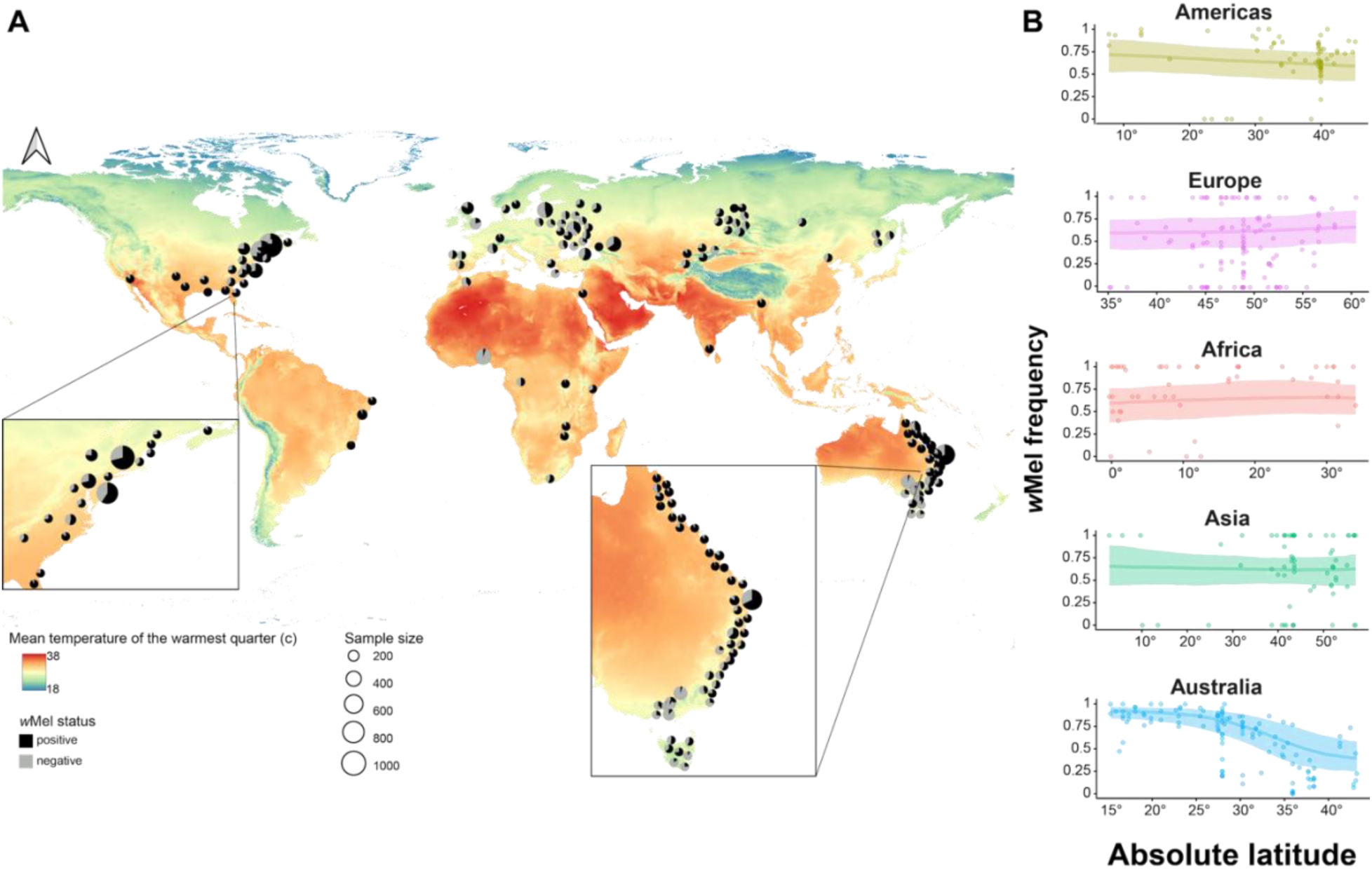
Global *w*Mel frequency variation across 248 locations, with a latitudinal cline restricted to Australia. **(A)** Global distribution of *w*Mel frequency across locations sampled between 1982 and 2024. Pie charts show the proportion of *w*Mel-positive (black) and *w*Mel-negative (grey) individuals at each location, with frequencies collapsed across temporal samples for presentation based on the general temporal stability reported by Kriesner et al. (2016) and confirmed here. Pie chart diameter is scaled to sample size; only locations with more than 10 individuals are shown. The base map reflects WorldClim BIO10 values (mean temperature of the warmest quarter, °C). Insets show eastern Australia and eastern North America. A latitudinal cline is evident in eastern Australia, with higher *w*Mel frequencies in tropical locations declining toward temperate latitudes (Kriesner et al. 2016); no comparable gradient is apparent in North America despite similar latitudinal coverage. Data sources: Kriesner et al. (2016), Soni et al. (2017), Рощина et al. (2018), Bykov et al. (2019), Gora et al. (2020), Cogni et al. (2021), Singhal and Mohanty (2025), our sampling from Linvilla, PA and Lohr’s, MD and genomic screening. A complete list of locations and sample sizes is in the SI Dataset. **(B)** Continent-specific relationships between absolute latitude and *w*Mel frequency from the Bayesian beta-binomial model with continent-specific spline terms for latitude. Lines show posterior mean predicted *w*Mel frequency as a function of absolute latitude (degrees) for each continent; shaded bands indicate 95% credible intervals. Points represent observed *w*Mel frequencies for individual location–time point observations. Australia was the only continent that showed a clearly non-linear decline in *w*Mel frequency with latitude (spline SD: 3.12 [95% CI: 1.36–6.25]); no latitudinal pattern was detectable for the remaining four continents.

A continent-specific spline model fit to the Edata revealed no consistent latitudinal cline across continents. Australia showed a clear non-linear decline in *w*Mel frequency with latitude (spline SD: 3.12 [95% CI: 1.36–6.25]; **Fig. 3B**); no latitudinal pattern was detectable for the remaining four continents (Africa: 0.75 [0.02–2.67]; Americas: 0.83 [0.03–2.69]; Asia: 0.58 [0.02–2.21]; Europe: 0.88 [0.03–3.04]), despite sampling spanning 35°–60°N in Europe. Region-specific Bayesian models confirmed the Australian cline on both datasets (Kdata spline SD: 1.91 [0.86–3.78]; Edata: 2.04 [0.92–4.08]) and the distinction between tropical Australia, where latitude had no detectable effect (Edata: 0.02 [−0.15, 0.19]), and temperate Australia, where *w*Mel frequency declined with latitude (Edata: −0.17 [−0.24, −0.11]). The significant cline below 38°N in North America detected by GLM (Kdata: b = −0.221, P < 0.001) was not supported by the Bayesian models (Edata: −0.13 [−0.50, 0.23]), reflecting overdispersion and non-independence that the GLM framework does not accommodate. The model identified substantial among-location variation in *w*Mel frequency (location SD: 0.50 [95% CI: 0.27–0.73]) that exceeded temporal variation within locations (SD: 0.28 [95% CI: 0.02–0.57]), indicating that spatial environmental differences are the primary axis of *w*Mel frequency variation globally. *w*Mel frequencies also differed among continents after accounting for latitude (continent SD: 0.62 [95% CI: 0.17–1.51]).

#### Bioclimatic predictors resolve the continent-level frequency structure that latitude cannot

Our bioclimatic model identified two global precipitation predictors (BIO15, precipitation seasonality; BIO17, precipitation of the driest quarter), one winter temperature predictor with an uncertain global effect (BIO6, minimum temperature of the coldest month), and one wet-season temperature predictor whose effect was continent-specific (BIO8, mean temperature of the wettest quarter) (**Table 1**). *w*Mel frequencies were higher where rainfall is more unevenly distributed across the year (BIO15; est.: 0.48 [95% CI: 0.24–0.71]; **Fig. 4A**) and where the driest quarter receives more rainfall (BIO17; est.: 0.43 [95% CI: 0.21–0.67]; **Fig. 4B**) — the two predictors with the strongest and most consistent global effects. *w*Mel frequencies were lower in locations with warmer winters than expected for their latitude (BIO6 residualized against latitude; est.: −0.30 [95% CI: −0.62 to 0.04]; **Fig. 4C**); the credible interval marginally overlapped zero in the bioclimatic model, but this association was clearly supported by the horseshoe model (est.: −0.44 [−0.78, −0.09]) and by the model without the BIO8 random slope (est.: −0.48 [−0.80, −0.19]). The attenuation occurs because BIO6 and the continent-specific BIO8 slope both capture aspects of seasonal temperature variation, so including the BIO8 slope absorbs some of the signal that BIO6 carried alone.

**Figure 4.**
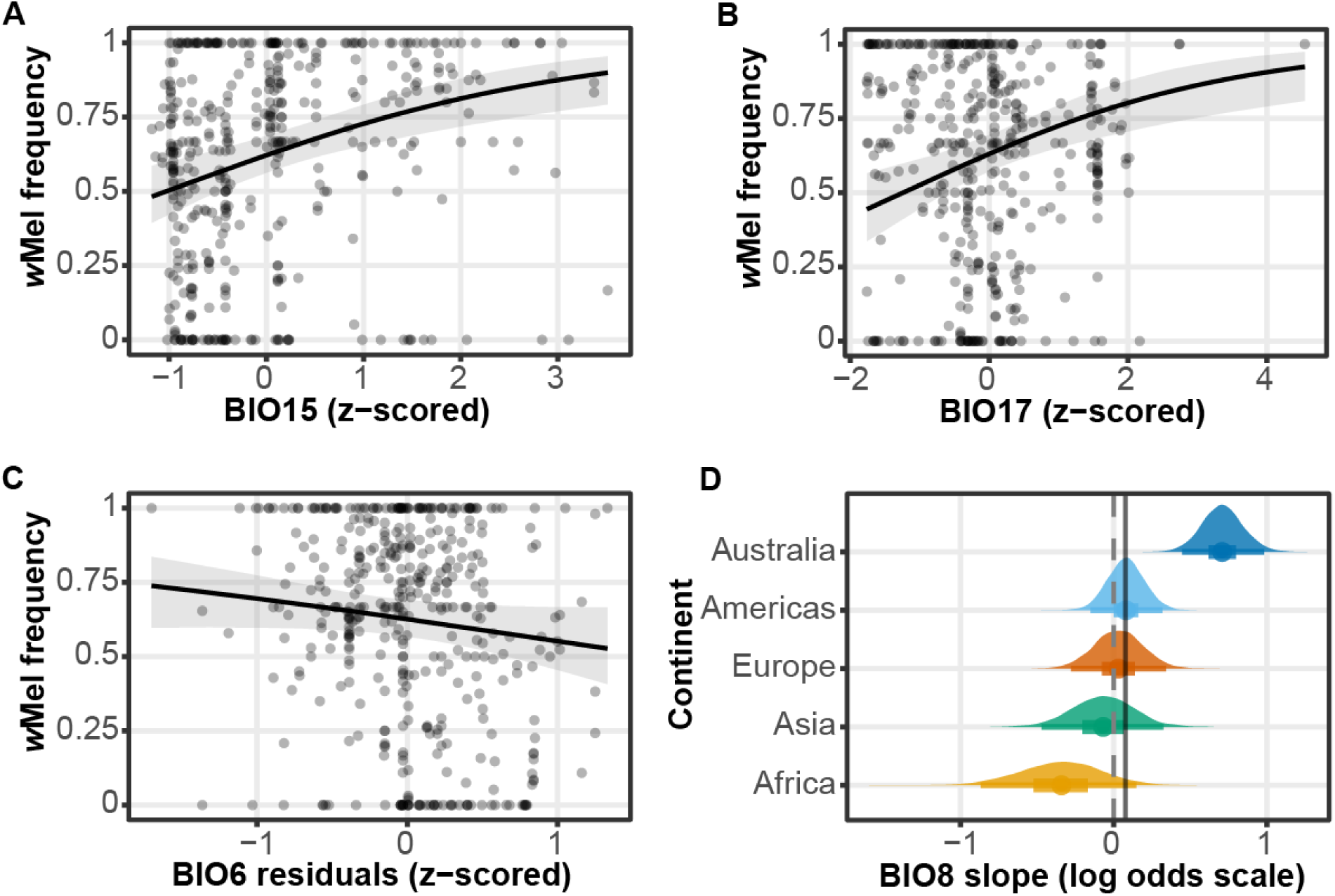
Precipitation regime and wet-season temperature predict *w*Mel frequency globally, with the temperature effect restricted to Australia. (A–C) Conditional effects of three global predictors: **(A**) scaled precipitation seasonality (BIO15), (**B**) scaled precipitation of the driest quarter (BIO17), and (**C**) residuals of scaled minimum temperature of the coldest month (BIO6). Predicted *w*Mel frequencies (y-axis) are shown as a function of each scaled bioclimatic variable (x-axis), with 95% credible intervals (shaded). Points represent observed *w*Mel frequencies for individual location-time point observations. Of these, BIO15 and BIO17 had credible intervals excluding zero; BIO6 was attenuated in the bioclimatic model because the continent-specific BIO8 slope absorbs overlapping seasonal temperature variation (see Results). (**D**) Continent-specific posterior distributions for the BIO8 (mean temperature of the wettest quarter) slope on the log-odds scale, estimated from the continent-specific random slope in the bioclimatic model. The dashed vertical line indicates zero (no effect); the solid vertical line indicates the global fixed effect estimate (0.08). Australia was the only continent with a 95% credible interval excluding zero, indicating a positive association between warm wet-season temperatures and *w*Mel frequency specific to Australia, where the wettest quarter coincides with summer and peak *D. melanogaster* reproductive activity. All predictions are from the bioclimatic model (see Methods).

**Table 1.** Bioclimatic model of global *w*Mel frequency: posterior summaries. All continuous predictors were standardized prior to model fitting to allow direct comparison of effect sizes. BIO6 (minimum temperature of the coldest month) was residualized against latitude before scaling to remove collinearity between the two predictors, so that the coefficient reflects thermal anomalies relative to latitude rather than raw temperatures. The intercept represents the expected *w*Mel frequency on the logit scale at mean values of all predictors. The fixed effect of BIO8 (mean temperature of the wettest quarter) represents the global average slope, while the continent-level random slope for BIO8 (sd continent: scaled BIO8) represents substantial among-continent variation, reflecting continent-dependent association between wet-season temperature and *w*Mel frequency. The model includes three levels of random intercepts: continent, location, and location:year, capturing variation in baseline *w*Mel frequency at the continental scale, among sampling locations, and among years within locations, respectively. Predictors were selected from 19 WorldClim bioclimatic variables using regularized horseshoe priors (see Methods) before refitting with standard priors. Posterior means, standard errors, and 95% credible intervals are reported for all fixed effects, random effect standard deviations, and the precision parameter (ϕ) of the beta-binomial distribution.

| Model parameter | Estimate | Estimated error | Lower 95% CI | Upper 95% CI |
| --- | --- | --- | --- | --- |
| Intercept | 0.53 | 0.13 | 0.28 | 0.81 |
| scaled residuals of BIO6 | -0.3 | 0.17 | -0.62 | 0.04 |
| scaled BIO8 | 0.08 | 0.25 | -0.44 | 0.56 |
| scaled BIO15 | 0.48 | 0.12 | 0.24 | 0.71 |
| scaled BIO17 | 0.43 | 0.12 | 0.21 | 0.67 |
| sd continent (Intercept) | 0.19 | 0.16 | 0.01 | 0.58 |
| sd continent (scaled BIO8) | 0.5 | 0.22 | 0.21 | 1.06 |
| sd location (Intercept) | 0.38 | 0.11 | 0.16 | 0.61 |
| sd location:year (Intercept) | 0.19 | 0.12 | 0.01 | 0.46 |
| $\phi$ | 5.53 | 0.74 | 4.27 | 7.17 |

Collectively, these predictors absorbed the continent-level variance that latitude left unexplained: continent random intercept SD – which measures residual among-continent variation in baseline *w*Mel frequency not explained by the fixed effects – dropped from 0.62 [95% CI: 0.17–1.51] in the latitude model to 0.19 [95% CI: 0.01–0.58] in the bioclimatic model, with all continent intercepts overlapping zero. Both models yielded marginal Bayesian R² = 0.83, but the bioclimatic model resolved the systematic geographic structure that the latitude model left behind. Leave-one-out cross-validation confirmed equivalent predictive performance (ΔELPD = −3.4, SE = 5.2; Sivula et al. 2025).

Mean temperature of the wettest quarter (BIO8) was retained by the horseshoe as a global predictor (est.: 0.21 [95% CI: 0.02–0.40]), but the bioclimatic model, which included a continent-specific random slope for BIO8, revealed that this signal was not globally uniform. This is expected for a predictor whose ecological meaning varies by continent: BIO8 measures wet-season warmth, but whether the wettest quarter aligns with the reproductive season differs fundamentally across continents (see Methods). The global BIO8 fixed effect was 0.08 [95% CI: −0.42–0.56], while the continent-specific random slope varied substantially (SD: 0.50 [95% CI: 0.21–1.06]; **Fig. 4D**). This variation was driven by Australia, which was the only continent with a BIO8 effect whose 95% credible interval excluded zero (continent-specific deviation: 0.63 [95% CI: 0.14–1.20]; total Australia slope: ∼0.71 on the log-odds scale). Europe showed no clear association (deviation: −0.04 [95% CI: −0.58–0.53]), and the Americas and Asia were similarly near zero (**Fig. 4D**). Africa trended negative (deviation: −0.43 [95% CI: −1.09–0.16]) but the credible interval overlapped zero. In Australia, the wettest quarter coincides with summer and peak *D. melanogaster* reproductive activity, where warm conditions promote maternal transmission fidelity (Hague et al. 2022). For example, in much of Europe the wettest quarter falls in winter when *D. melanogaster* populations are dormant or at very low densities, and in parts of North America and Asia peak rainfall occurs in spring or autumn, decoupling BIO8 from the period of active reproduction (see also **Fig. S4** for descriptive visualization of raw *w*Mel frequencies against warmest-quarter temperature).

#### wMel frequencies are broadly stable through time, but short-term and directional changes occur at specific sites

Independent of the bioclimatic associations described above, we quantified temporal *w*Mel frequency changes at 28 locations with three or more temporal observations, encompassing 16 locations with within-year sampling and 27 locations with multi-year sampling (**Table S15**). At short temporal scales, GLMs and G-tests agreed on significant variation at 4 of 16 locations (Lohr’s, Linvilla, Gold Coast, and Uman), while G-tests alone detected significant heterogeneity at 4 additional locations where GLMs either lacked power or could not be fit (**Table S15**). (G-tests detect any frequency heterogeneity among sampling periods, while GLMs test for a systematic directional trend and are more sensitive to sample size.) At long temporal scales, 2 of 27 locations showed significant multi-year variation by both tests (Cobram and Chernobyl), GLMs alone detected a significant positive trend at Hastings (b = 0.049, *P* = 0.018; *G* = 3.96, *P* = 0.138), and G-tests detected significant heterogeneity at 8 additional locations (**Table S15**). The majority of locations showed no significant directional trend by GLM, indicating that *w*Mel frequencies are broadly stable across years at most sites. That G-tests detected heterogeneity at more locations than GLMs is consistent with non-directional frequency fluctuation of the kind documented within seasons at our orchard sites and at Gold Coast. Modeling of stochastic *Wolbachia* dynamics under realistic *w*Mel parameters predicts that weak-CI strains are inherently prone to temporal fluctuation, but that stochastic dynamics alone cannot account for the largest observed frequency shifts (Graham et al. 2025). Whether such fluctuation is common but undetectable at less intensively sampled sites remains an open question.

### *w*Mel SNP-environment associations reflect lineage structure, not local adaptation

We tested whether *w*Mel allele frequencies were associated with latitude or bioclimatic variables across 339 individually sequenced genomes from 34 populations on five continents, using permutation-based genome-wide significance thresholds. Genome-wide permutation tests identified significant associations between *w*Mel allele frequencies and two environmental predictors: latitude (*P* < 0.001) and mean temperature of the wettest quarter (BIO8; *P* < 0.001). Of the 593 biallelic SNPs, 38 were significantly associated with latitude, and all 38 showed higher derived allele frequencies at temperate latitudes (**Table S16**; beta range: 0.97–2.50, all 95% CIs above zero). SnpEff annotations indicated 19 missense variants, 6 synonymous variants, and 13 intergenic or upstream/downstream modifiers among the 38 (**Table S16**); however, the ancestry corrections described below indicate that these annotations should be interpreted as properties of the lineage carrying the derived alleles rather than as evidence for locus-specific selection. The only bioclimatic association exceeding the permutation-based genome-wide threshold was a single synonymous SNP in *ruvB*, a gene involved in recombinational DNA repair, which was positively associated with BIO8 (beta = 3.24, 95% CI: 2.10–4.38).

Pairwise linkage disequilibrium was near-complete across the *w*Mel genome (median r² = 0.94; median |D′| = 1.0), with negligible decay with physical distance (Spearman ρ = −0.042 between r² and distance). Four-gamete violations occurred at 2.95% of SNP pairs. These violations were concentrated at large genomic distances, consistent with either low-level recombination or genotyping artifacts, but the near-complete LD across the full genome indicates that all 593 SNPs effectively share a single genealogy. A maximum-likelihood phylogeny of the 339 individually sequenced *w*Mel genomes, rooted on sister *w*Zts found in *Zaprionus tsacasi* (Shropshire et al. 2026), resolved five genomes from Montpellier, France and Azeitão, Portugal as a monophyletic clade diverging after clade VI (*w*MelCS-like; Richardson et al. 2012) but before the main *w*Mel radiation encompassing clades I–IV and VIII, which contains most global diversity (**Fig. 5A**). These five genomes each carried ≥36 of the 38 latitude-associated derived alleles (four carried all 38; one Azeitão genome retained the ancestral state at two positions). Thirty-five of the 38 were absent from all other lineages, consistent with mutations that accumulated on the branch leading to this clade. The remaining three were shared with four clade VI genomes – two from Raleigh, North Carolina; one from Sorrell, Tasmania; and one from Cooktown, Queensland – which each carried only these three derived alleles at the same position (805011, 812321, and 1206452) and none of the other 35. Because clades V and VI are the earliest-diverging *w*Mel lineages (**Fig. 5A**), these three alleles most likely represent the ancestral *w*Mel state, retained in clades V and VI and lost in the lineage leading to clades I–IV and VIII. These alleles are classified as derived relative to the *w*Zts outgroup used for polarization, but are ancestral within *w*Mel based on their presence in the two earliest-diverging clades. The remaining 330 genomes across all continents carried none of the 38 derived alleles. Derived allele frequencies at the 38 SNPs were concentrated almost entirely in Europe (mean AF 0.22, median 0.17), with negligible frequencies in Australia (0.016), the Americas (0.002), and Africa (<0.001).

**Figure 5.**
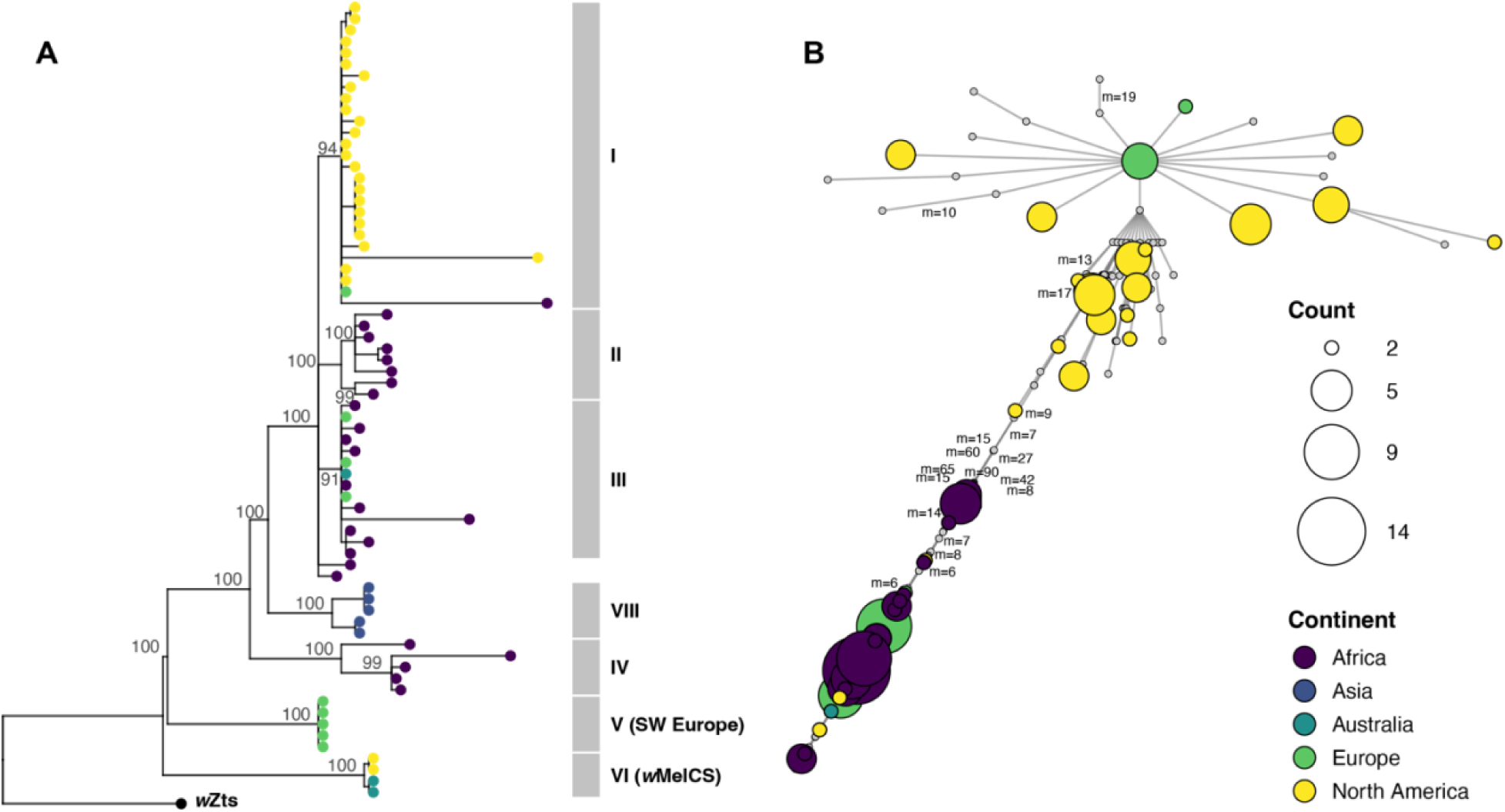
*w*Mel genomic diversity reflects geographically structured lineage history, not independent allele-environment associations. **(A)** Maximum-likelihood phylogeny of a representative subset (*N* = 70) of 339 individually sequenced *w*Mel genomes (pruned from the full tree for presentation; see Fig. S6 for the complete tree), based on 728 single-copy genes (736,557 bp) and rooted on *w*Zts from the sister species *Z. tsacasi* (Shropshire et al. 2026). Pruning retained all members of clades IV, V, VI, and VIII in full, and subsampled clades I–III proportionally; topology is identical between the pruned and full trees. Tip circles are colored by continent of origin. Numbers at nodes indicate bootstrap support (%). Grey bars denote cytoplasmic clade designations following Richardson et al. (2012). Clades I, II, III, IV, VI and VIII are confirmed by reference genomes included in the analysis. Clades I, II, and III form an unresolved polytomy in our phylogeny, whereas Richardson et al. resolved a short internal branch separating them. Five genomes (Montpellier, France and Azeitão, Portugal) form a monophyletic clade diverging after clade VI but before clades IV, VIII and I–III (SW European lineage; assigned to clade V by mitochondrial haplotype affinity, see Methods). All five genomes of the SW European lineage carry ≥36 of the 38 derived alleles, four clade VI genomes (two from Raleigh, NC, USA; one from Sorrell, Tasmania; one from Cooktown, Queensland) carry exactly three, and the remaining 330 genomes carry none. **(B)** Haplotype network constructed from all 593 biallelic SNPs using minimum-spanning network algorithm. Node size is proportional to the number of genomes sharing a haplotype, node color indicates continent of origin, and small open circles represent unsampled intermediate haplotypes. Edge labels indicate the number of mutational steps (*m*); unlabeled edges represent single steps. African genomes are concentrated in one cluster and North American genomes in the other, connected through intermediate haplotypes; European genomes span both clusters, with the five clade V genomes embedded within the African cluster.

A haplotype network constructed from all 593 SNP positions confirmed geographically structured *w*Mel diversity, with African genomes concentrated in one cluster and North American genomes in the other, connected through intermediate haplotypes (**Fig. 5B**). European genomes were distributed across both clusters, and geographic structure in the network broadly reflected continental origin rather than latitude.

Even within non-tropical populations, derived alleles at the 38 SNPs occurred at low frequency (∼2.6%), and 35 of 38 alleles were absent from tropical populations. We repeated the genome scan with covariates that correct for progressively finer scales of *w*Mel lineage structure. Replacing hemisphere with an Africa/non-Africa binary covariate eliminated all 38 associations (0/38 significant; 10,000 permutations, LRT threshold: 38.88). A Europe/non-Europe covariate retained 8 of the 38 SNPs (LRT threshold: 24.71), but a first principal component (PC1) computed from the 555 non-latitude-associated SNPs eliminated all but a single intergenic SNP near *mnmA* (1/38; LRT 34.93, threshold 33.14; 10,000 permutations). This PC1 was highly correlated with a PC1 computed from all 593 SNPs (*r* = 0.995), confirming that *w*Mel lineage structure is genome-wide rather than driven by the 38 latitude-associated SNPs. Scans restricted to Europe and North America separately yielded no significant associations (0/38 for both), confirming the absence of signal within continents. Eight of the 38 latitude-associated SNPs initially overlapped with the 43 candidates (Hague et al. 2022) identified using pool-seq data; four of those eight survived under the Europe/non-Europe correction but were eliminated by PC1, indicating that the overlap reflects shared sensitivity to *w*Mel lineage structure rather than independent confirmation of environment–allele associations.

We next tested whether allele frequency variation in the *w*Mel genome differs between nonsynonymous and synonymous sites or is concentrated in the WO prophage region, focusing on the 38 latitude-associated SNPs and the full set of 593 biallelic SNPs. Latitude-associated SNPs were not enriched in the WO prophage region relative to the background SNP set (Fisher’s exact test, OR = 1.34, *P* = 0.50). Separately, eight of the 38 fell within or adjacent to prophage-associated genes by SnpEff annotation, but only a *pleD* family response regulator missense variant was physically located within prophage gene boundaries (**Table S16**). Nonsynonymous sites were not enriched among the 38 SNPs relative to background (Fisher’s exact test, OR = 0.84, *P* = 0.82), and the folded site-frequency spectra of nonsynonymous and synonymous sites across all 593 SNPs were indistinguishable (Kolmogorov-Smirnov test, *P* = 0.67). A skew in the folded site-frequency spectrum toward rare alleles (**Fig. S7A**) produced negative Tajima’s *D* (−1.84), but a heterogeneity test (Hahn et al. 2002; 10,000 coalescent simulations, *N* = 335) confirmed the skew did not differ between synonymous (−1.96) and nonsynonymous (−1.75) sites (*P* = 0.58). Unfolding the site-frequency spectrum with *w*Ri as an outgroup (∼7.5 MY divergence) showed that derived synonymous alleles spanned a wider frequency range than nonsynonymous alleles (IQR: 0.702 vs. 0.319, **Fig. S7B**), but the overall distributions did not differ significantly (Kolmogorov-Smirnov test, *P* = 0.65), and power was limited with 65 synonymous sites. Allele frequency variation in the *w*Mel genome is thus not concentrated in nonsynonymous changes or in the prophage region, consistent with the near-complete linkage across the *w*Mel genome, in which all sites effectively share a single genealogy.

## Discussion

Climatic conditions – not latitude per se – predict global *w*Mel frequency variation in *D. melanogaster*. At the orchard scale, *w*Mel frequencies shifted by up to 0.33 between weekly samples, cages followed different frequency trajectories, and frequencies were consistently higher in June than November across seven years of sampling at Linvilla, PA. At the global scale, analysis of 248 locations across 42 years confirmed substantial *w*Mel frequency variation that latitude could not explain outside of eastern Australia (Kriesner et al. 2016). Our bioclimatic model matched the latitude model in overall explanatory power, while resolving the systematic continent-level structure that latitude left behind. The model identified precipitation regime as the primary global axis and implicated wet-season temperature as an Australia-specific contributor, where unlike other continents the wettest quarter coincides with summer and peak host reproduction. An early-diverging *w*Mel lineage restricted to southwestern Europe generated the allele frequency structure we detected – an association with latitude that dissolved under corrections for *w*Mel lineage structure – and this genomic variation was not preferentially concentrated in protein-coding changes. Below we discuss these findings in the context of global *w*Mel spread and the mechanisms governing its frequency dynamics.

### Temperature drives *w*Mel frequency variation at the orchard scale

Hoffmann et al. (1998) documented marked *w*Mel frequency fluctuations between monthly collections at Gold Coast, Australia, with frequencies dropping from ∼0.92 to ∼0.25 across a single monthly interval. We observed changes of comparable magnitude on even shorter weekly timescales, with *w*Mel frequency consistently peaking at intermediate temperatures and declining at both thermal extremes across three independent locations. This non-linear association is consistent with temperature-dependent maternal transmission, where cool rearing temperatures reduce *w*Mel abundance at the posterior pole of developing oocytes and impair transmission fidelity in the laboratory (Hague et al. 2022). Effects of temperatures above 28°C on *w*Mel transmission are unknown in *D. melanogaster*, but the frequency decline we observed at higher temperatures parallels heat effects on *w*Mel transinfections in *Ae. aegypti* (Ross et al. 2017; Mancini et al. 2021). Microhabitat-level differences in sun exposure add further support: because all experimental cages were founded simultaneously from the same outbred stock at statistically indistinguishable starting frequencies, and because subsequent divergence clustered by shade status rather than randomly among cages, the opposing trajectories implicate thermal microenvironment rather than founder effects or genetic drift. Given that *w*Mel CI is weak and declines rapidly with male age (Shropshire et al. 2021), variable CI strength is unlikely to contribute substantially to the frequency variation we observed (Graham et al. 2025). Confirming the role of temperature-dependent maternal transmission will require field estimates of transmission fidelity and host fitness effects paired with concurrent frequency tracking.

The long-term Linvilla data confirmed a seasonal signal, with *w*Mel frequencies significantly higher on average in June than November across seven years of sampling. Mean temperatures at Linvilla during these periods differed substantially: 22.1°C in June, near the thermal optimum we identified at the orchard scale, versus 13.7°C in October and 7.6°C in November, well below the temperatures that reduced *w*Mel transmission fidelity in the laboratory (Hague et al. 2022). Seasonal variation parallels the temporal instability Hoffmann et al. (1998) documented at Gold Coast, Australia, while the long-term stability across years mirrors the general temporal stability of the eastern Australian cline (Kriesner et al. 2016). Linvilla lies above the 38°N threshold at which Kriesner et al. (2016) suggested *D. melanogaster* populations may not persist outdoors year-round. However, repeatable seasonal allele frequency oscillations and phenotypic signatures of winter selection at Linvilla support that a resident population persists through winter bottlenecks (Bergland et al. 2014; Behrman et al. 2015), consistent with the elevated overwintering F_st_ relative to within-season F_st_ observed at comparable latitudes (Nunez et al. 2025). The seasonal *w*Mel frequency difference we observe may therefore reflect winter selection against *Wolbachia*-carrying females during the bottleneck (Kriesner et al. 2016), as well as within-season temperature effects on maternal transmission fidelity (Hague et al. 2022; Hague et al. 2024) and potentially other parameters (*e.g.,* CI; Clancy and Hoffmann 1998; Nasehi et al. 2022; Bagchi et al. 2026).

### Latitude as a continent-specific proxy for *w*Mel frequency variation

Our Bayesian analysis with continent-specific splines confirmed two of Kriesner et al.’s findings simultaneously. First, Australia was the only continent showing a clear non-linear decline in *w*Mel frequency with latitude (spline SD: 3.12 [95% CI: 1.36–6.25]), the pattern Kriesner et al. approximated as a linear cline. Second, the model estimated no latitudinal relationship for Africa, Asia, or Europe, shrinking spline SDs toward zero when the data provided insufficient evidence for a pattern. Kriesner et al. reached similar conclusions, finding no latitudinal pattern in Eurasian data and noting highly variable frequencies in equatorial Africa—from over 96% *w*Mel frequency in Rwanda to less than 10% in Ghana. They interpreted their findings through the Hoffmann et al. (1990) equilibrium framework, proposing that fitness costs during reproductive dormancy could reduce frequencies at high latitudes, while weak CI from young males may elevate tropical frequencies. Dormancy plausibly operates in eastern Australia where *D. melanogaster* populations persist year-round across a latitudinal gradient in winter severity, but provides less insight into why frequencies vary among continents or among equatorial populations. Consistent with this framework, the temperate Hastings site in the southern part of the cline showed a significant positive directional trend in *w*Mel frequency across years (b = 0.049, *P* = 0.018), matching Kriesner et al.’s prediction that warming winters should increase frequencies at high-latitude sites.

Kriesner et al. also reported a shallower latitudinal cline in eastern North America, finding a significant negative association after excluding locations above 38°N where *D. melanogaster* populations may not persist through winter. Our GLMs replicated their significant negative association below 38°N, but the Bayesian model fit to the same data did not. DHARMa results confirmed that this discrepancy reflects overdispersion and non-independence that the GLMs ignore, but the beta-binomial likelihood and location random intercepts account for (**Table S14**). The Bayesian model also failed to detect a latitudinal signal across all locations in eastern North America, which has a plausible demographic explanation. Kriesner et al. proposed that high-latitude North American populations may be re-colonized each spring from local refugia or by human-mediated transport, rather than maintained by overwintering adults (see Wilfert and Jiggins 2014). If so, observed *w*Mel frequencies at high latitudes would reflect source-population composition rather than local transmission-selection equilibrium. Whether dormancy costs contribute to *w*Mel frequency at a given location depends on whether populations persist locally across seasons. Some North American populations near and above 38°N clearly do (Bergland et al. 2014; Behrman et al. 2015; Nunez et al. 2025), while others at higher latitudes may not—a distinction that varies among locations and that the Bayesian model accommodates through location-level deviations rather than an imposed latitude threshold.

Identifying which environmental conditions modulate *w*Mel transmission fidelity and effects on hosts is central, because even small geographic differences could generate the among-population variation in *w*Mel frequency that latitude leaves unexplained (Kriesner et al. 2016). Temperature clearly affects *w*Mel transmission fidelity (Hague et al. 2022; Hague et al. 2024), but *w*Mel CI strength was indistinguishable between warm (26°C) and cool (18°C) laboratory temperatures (Bagchi et al. 2026). CI declines rapidly with male age in *D. melanogaster* (Reynolds and Hoffmann 2002; Yamada et al. 2007; Shropshire et al. 2021), and as noted by Kriesner et al., geographic variation in the age distribution of mating males could contribute to *w*Mel frequency variation. Even with plausible CI contributions, substantial positive fitness effects are required to explain very high *w*Mel frequencies (*e.g.*, ∼90%) (Kriesner et al. 2016). While variable fecundity effects (Olsen et al. 2001; Fry et al. 2004; Gruntenko et al. 2017; Serga et al. 2021), pathogen protection (Teixeira et al. 2008; Osborne et al. 2009; Martinez et al. 2014; Shi et al. 2018), and nutritional benefits remain candidates (Brownlie et al. 2009), the specific fitness effects contributing to *w*Mel frequency and sensitivity to the environment are mostly unresolved. The bioclimatic predictors we examine below do not resolve these mechanisms, but they identify the environmental axes along which *w*Mel frequency varies globally.

### Bioclimatic predictors resolve what latitude cannot

Precipitation seasonality (BIO15) and driest-quarter precipitation (BIO17) carried the strongest global effects, with credible intervals excluding zero. Although these variables are negatively correlated climatically – greater rainfall seasonality produces drier dry quarters – both positively predicted *w*Mel frequency, indicating they capture distinct biological mechanisms. These variables likely capture host demographic and nutritional conditions rather than direct effects on *Wolbachia*: high precipitation seasonality characterizes environments with marked wet-dry contrasts, where seasonal resource pulses drive large increases in host population size (Behrman et al. 2015; Mansourian et al. 2018), and higher driest-quarter precipitation may sustain host access to fruit resources during otherwise resource-limited periods. Serbus et al. (2015) demonstrated that host diet affects *Wolbachia* titer in *D. melanogaster* oocytes through TOR and insulin signaling — the same pathways that regulate host fecundity and reproductive investment in response to resource availability (Drummond-Barbosa and Spradling 2001; LaFever and Drummond-Barbosa 2005; LaFever et al. 2010; Paaby et al. 2010; Flatt 2020). Nutritional variation could therefore modulate *w*Mel frequency simultaneously through its effects on both maternal transmission fidelity and host fitness.

BIO8 showed no consistent global effect, as expected for a predictor whose ecological meaning varies by continent, but the Australia-specific BIO8 effect helps explain why latitude predicts *w*Mel frequency in eastern Australia. The wettest quarter in Australia coincides with summer, a period of peak *D. melanogaster* reproductive activity and when warm conditions plausibly promote maternal transmission fidelity (Hague et al. 2022). Hence, latitude in Australia correlates simultaneously with temperature during reproduction, winter severity, and the seasonal alignment of peak rainfall with reproductive activity. In contrast, the wettest quarter falls in winter in much of Europe when *D. melanogaster* populations are dormant or at very low densities, while in parts of North America and Asia, peak rainfall occurs in spring or autumn, decoupling wet-season warmth from peak reproduction. This seasonal misalignment plausibly explains why BIO8 contributes substantially to predicting *w*Mel frequency within Australia but explains little variation globally.

Although winter severity contributes to latitude’s effectiveness in Australia, the global winter temperature predictor (BIO6) proved harder to interpret. Because we statistically removed the correlation between BIO6 and latitude before entering BIO6 into the model, its coefficient isolates winter thermal anomalies from the latitudinal gradient in winter severity itself; raw BIO6 and latitude are too collinear to include in the same model (see Methods). The dormancy effect Kriesner et al. documented operates along the latitudinal gradient – colder at higher latitudes – and is therefore absorbed by the latitude term before BIO6 residuals enter the model. What the negative residualized BIO6 coefficient captures is the additional variation: among locations at the same latitude, those with anomalously mild winters have lower *w*Mel frequencies. This is consistent with our observation at Linvilla that seasonal frequency changes track temperature independently of dormancy. Locations that are anomalously warm in winter for their latitude may differ from colder counterparts in growing season length, resource dynamics, or host demography in ways that independently affect *w*Mel frequency. Disentangling these possibilities will require field data pairing *w*Mel frequency with direct measurements of host nutritional status and dormancy duration, while also quantifying micro-habitat environmental variation that will more accurately reflect the conditions experienced by flies in nature.

Taken together, the precipitation and temperature predictors suggest that *w*Mel frequencies reflect local ecological conditions – including resource availability, host demography, and the seasonal alignment of favorable conditions with reproduction – rather than geography per se.

### Lineage structure, not environment-based selection, underlies *w*Mel genomic variation

Recombination between *Wolbachia* genomes has been documented across divergent strains (Jiggins et al. 2001; Werren and Bartos 2001; Baldo et al. 2006), and its presence or absence within *w*Mel determines whether geographic patterns in allele frequencies reflect independent changes at individual loci or the sorting of lineages. While single-copy loci show little evidence for recombination or horizontal allele acquisition across *w*Mel-like variants spanning millions of years of host divergence (Shropshire et al. 2026), WO prophage and CI loci show evidence of gain and loss on shorter timescales, indicating that genomic compartments have distinct evolutionary histories. The near-complete LD we observed indicates that all 593 SNPs effectively share a single genealogy within *D. melanogaster*.

Our ancestry corrections confirmed that the 38 latitude-associated SNPs mark this genealogy’s geographic structure rather than independent environment-allele associations. The signal dissolved completely within Europe and within North America, and correcting for the first principal component of *w*Mel variation eliminated it genome-wide. The five southwestern European genomes that carry the derived alleles form a monophyletic clade diverging after clade VI – the *w*MelCS-like remnant that Richardson et al. (2012) documented as evidence for incomplete replacement of *w*MelCS by *w*Mel – but before clades IV, VIII, and I–III and the main radiation encompassing most global diversity. Previously characterized from a single Portuguese genome and mitochondrial haplotypes of two uninfected French strains (Richardson et al. 2012; Versace et al. 2014), the clade V lineage carries 35 private derived alleles that accumulated on the branch leading to it, and whose geographic restriction to southwestern Europe generated the latitude association we detected. The parallel with clade VI is direct: both represent lineages that diverged early in *w*Mel’s ∼8,000-year spread and persisted in restricted ranges while derived haplotypes radiated globally. Versace et al. (2014) demonstrated that cold experimental conditions reproducibly shifted clade composition in favor of clade V at the expense of other clades, in a response attributable to *w*Mel rather than to host nuclear background or mtDNA.

Eight of the 38 latitude-associated SNPs overlapped with the 43 candidates Hague et al. (2022) identified by comparing tropical and temperate Australian *w*Mel variants that differed in maternal transmission fidelity. We found no significant association between *wspB* allele frequency and latitude or any bioclimatic predictor, despite *wspB* being their strongest candidate. Because their tropical variant belongs to clade I and their temperate variant to clade III, both studies effectively recovered SNPs distinguishing early branches of the *w*Mel genealogy rather than loci under environment-driven selection. Clade-level fitness differences may well exist – the Versace et al. (2014) cold-evolution result suggests they do – but genome scans in symbionts like *w*Mel, where near-complete linkage ties all loci to a single genealogy, cannot distinguish selection at individual loci from the sorting of lineages that differ at many sites simultaneously.

### Conclusions

Our results establish that *w*Mel frequencies can vary on the order of weeks and, in our experimental orchard, across meters of microhabitat, tracking temperature in a manner consistent with known effects on maternal transmission fidelity (Hague et al. 2022). At the global scale, precipitation and temperature predictors resolve the continent-level frequency structure that latitude cannot, implicating host nutrition and population dynamics alongside temperature as plausible drivers of *w*Mel frequency variation. These results support a model in which long-term climatic conditions – including precipitation regime and thermal environment – set equilibrium *w*Mel frequencies that differ among populations, while short-term temperature fluctuations transiently perturb frequencies around those equilibria, as we documented at the orchard scale. This accommodates the general temporal stability Kriesner et al. (2016) observed across the Australian cline, with transient perturbations (*e.g.,* Hoffmann et al. 1998) producing the rapid temporal variation we documented here. We identified 35 private derived alleles in five clade V *w*Mel genomes from southwestern Europe—an early-diverging lineage whose geographic restriction generated the latitude associations we detected. This underscores that *w*Mel variant biogeography remains incompletely catalogued despite decades of study.

Connecting the environmental gradients we identify to the specific effects governing *w*Mel frequency in natural and transinfected systems will require field experiments that jointly estimate maternal transmission, components of host fitness, and CI under realistic conditions. Identifying targets of selection in *w*Mel genomes, where near-complete linkage ties all loci to a single genealogy, will require genomic analysis of populations sampled through time that can distinguish allele frequency change from static lineage structure. Moreover, applications involving population replacement rely on maintaining pathogen-blocking *w*Mel transinfections at high frequencies in *Ae. aegypti* populations to reduce dengue incidence (Hoffmann et al. 2011; Utarini et al. 2021; Lenharo 2023; Velez et al. 2023; de Morais Batista et al. 2026), and the efficacy of these programs depends on local environmental conditions (Ross et al. 2017; Ross, Ritchie, et al. 2019; Gesto et al. 2021; Hien et al. 2022; Hoffmann and Cooper 2025). The bioclimatic approach we implemented may help anticipate where *Wolbachia* frequencies are most vulnerable to environmental disruption, as environmental change alters both temperature and precipitation regimes.

## Materials and Methods

### Establishing and sampling cages in an experimental orchard

Our work builds on more than a decade of studies at an experimental orchard in Pennsylvania (Rudman et al. 2022). To initiate our experimental cage populations, we first recombined and expanded 77 inbred fly lines derived from wild-caught individuals collected from Linvilla, PA in 2012 (Behrman et al. 2015) to create a genetically diverse outbred founder population. In June of 2023, we used 500 males and 500 females from the founder population to establish each independent, outdoor cage population (*N =* 12) in our experimental orchard.

Each cage population inhabited a 2m x 2m x 2m meshed enclosure encompassing a dwarf peach tree, with clover planted as ground cover, mimicking a natural insect and microbial community. We provided the flies with Spradling/Bloomington cornmeal molasses-based medium on aluminum loaf pans added every other day during population establishment and from July 7, through October 14, 2023, which comprised the period of our 14-week experiment. The loaf pans were covered with a mesh lid after two days of oviposition and placed in a 0.3m x 0.3m x 0.3m meshed eclosion chamber inside their respective 2m x 2m x 2m enclosure.

We used an aspirator to collect 100-150 flies from each eclosion chamber within each of the 12 larger enclosures each week from July to October. Remaining flies in each eclosion chamber were then released into their respective larger enclosure. While the population in the larger enclosure consisted of flies of different ages, flies in the eclosion chamber were not older than 5 days post eclosion. We placed HOBO loggers in each of the 12 enclosures for the duration of the experiment. However, data was available for the entire duration of the experiment from four loggers, placed in enclosures E01, E04, E07, and E08. We calculated weekly mean and weekly average maximum temperature independently for each logger. We then tested if weekly mean and weekly average maximum temperature among HOBO loggers differed significantly using the statistical methods described below.

### Sampling of natural orchards populations

We sampled *D. melanogaster* from two natural orchards at times that overlapped with our sampling of the experimental UPenn orchard. This included weekly collections of *D. melanogaster* from July 10 to August 23, 2023, at Linvilla, PA and at Lohr’s Orchard, MD, which corresponded to weeks 2–7 described above for the experimental orchard. We also sampled Lohr’s Orchard from September 9 to October 4, 2023, which corresponded to weeks 11–14 described above. We aspirated flies directly from fruit substrates (*i.e.,* cherries, peaches, and apples) into vials containing Spradling/Bloomington cornmeal molasses-based medium. We established isofemale lines by separating single gravid females into individual vials with cornmeal molasses-based medium. To distinguish *D. melanogaster* from *D. simulans* lines, we examined the genital arch of F_1_ male offspring that emerged from each vial. Lines identified as *D. simulans* were excluded from our study. We obtained daily temperature data (NSRDB) for each location for the duration of the sampling from the USA and Americas (2018-2024) database from https://nsrdb.nrel.gov/data-viewer at 30- and 60-minute intervals.

We complemented our intensive weekly sampling with prior unpublished data from seasonal sampling of Linvilla from 2009 to 2015. Flies were collected using a combination of baited traps and aspirating flies from fruit substrates. Flies were sorted by species using light CO_2_ and gravid female were used to establish isofemale lines from each collection. *D. melanogaster* and *D. simulans* were distinguished using the method described above. Flies were maintained in standard laboratory culture (25°C 12:12 L:D) on vials containing Spradling/Bloomington cornmeal molasses-based food. After the isofemale lines were established, the F_1_ progeny were preserved for each line in 95% ethanol.

### Screening orchard populations for *w*Mel

For our 2023 sampling of each cage population and both natural orchards, we screened a minimum of 10 females from each available sample (**Table S1–S3, S5–S6**). We extracted DNA from individual flies in 96-well plates using a standard “Squish” buffer extraction protocol (10 mL Tris-HCl [1M], 0.0372 g EDTA, 0.1461 g NaCl, 90 mL dH20, followed by 150 mL Proteinase K after autoclaving) (Gloor et al. 1993). Each plate included DNA extractions from flies of known positive and negative *Wolbachia* status as controls. For our seasonal Linvilla sampling, we screened lines (*N* = 30) from each collection by pooling five females from each line for DNA extractions using Squish buffer. We used standard polymerase chain reaction (PCR) to type each sample for *w*Mel, using the *wsp* gene primer for *Wolbachia* along with the arthropod specific *28S* rDNA primer as positive control (Cooper and Shropshire 2024).

### Analysis of global *w*Mel data

#### Analyzing global wMel frequency

To further evaluate the potential for temporal and spatial *w*Mel frequency variation, we collated publicly available *w*Mel frequency estimates as of August 23, 2025. These included the data analyzed by Kriesner et al. (2016), our frequency estimates for the two natural orchards described above and estimates from six studies published after 2016 (Soni et al. 2017; Рощина et al. 2018; Bykov et al. 2019; Gora et al. 2020; Cogni et al. 2021; Singhal and Mohanty 2025). We additionally screened two publicly available bioprojects consisting of individually sequenced *D. melanogaster* genomes that were not represented in Kriesner et al. (2016), using a two-stage process to identify *w*Mel-positive samples. First, we used Magic-BLAST (Boratyn et al. 2019) to compare the raw reads to *w*Mel *fbpA*, *ftZ*, *groE*, *coxA* and *hcpA* sequences, retaining only those reads with matches >100bp in length, >98% similarity to the reference genes, and at least five sequence reads for each gene (Pascar and Chandler 2018). Second, we generated reference-guided whole-genome alignments for all samples, trimming raw reads with *fastp* (Chen et al. 2018) and mapping against the reference *w*Mel genome (GCF_000008025.1) using bwa-mem2 (Li 2013) with default parameters. We used SAMtools (Li et al. 2009) to quantify genome-wide means for read depth and coverage, retaining samples with >20x mean read depth and >5x coverage as *w*Mel positive. While many pooled-sequenced samples exist (*e.g.,* Kapun et al. 2021), we did not include these data because the total number of *Wolbachia*-positive individuals in each pool cannot be ascertained. We identified duplicated observations in the Kriesner et al. (2016) dataset (see Supplemental Materials) and removed them prior to analysis unless explicitly noted.

#### Calling SNPs in wMel genomes

Allele frequencies among *w*Mel-positive reads can be estimated directly from read counts at each position, conditional on the sample having passed our *w*Mel screening criteria. Hence we used the individually sequenced samples that passed our criteria above (*N* = 339 sequences), for all individually sequenced *D. melanogaster* samples (*N* = 460) from 34 natural populations and additional pooled-sequencing data (*N* = 383 pools) from 136 natural populations, to analyze *w*Mel allele frequencies. Our final dataset consisted of 722 samples combined from individually and pooled sequence reads. We identified *w*Mel-positive pools using the criteria above, except for only retaining samples with >40x mean read depth and >20x mean coverage. This enabled us to account for the increased variance in allele representation in pooled-sequenced samples. Samples meeting our criteria were deduplicated using picardtools (Tools 2019) and sorted by genome coordinates using SAMtools. We used bcftools *mpileup* (Danecek et al. 2021) to generate genotype likelihoods from aligned reads and bcftools *call* to call variants with a minimum mapping quality of 20, base quality of 30 and a maximum read depth to 200. To reduce false SNP positives, we applied additional filters only on individually sequenced samples to identify a true SNP set. We specifically excluded pooled sequenced reads for SNP identification as unknown and variable *w*Mel prevalence across pooled samples can lead to inflation of read depth as SNP. We retained only those SNPs with an alternate allele count ≥ 2 and an alternate allele count ≤ AN − 2, effectively excluding singletons and nearly fixed sites, a root mean mapping quality >40, and a variant call quality score >30, resulting in 593 biallelic SNPs. We ran bcftools *mpileup* and bcftools *call* again on all QC passed samples for these 593 variants. We calculated population level allele frequency estimates using bcftools *fill-tags* and custom bash scripts (https://github.com/nitinra/pa_experiment.git) <u>based on allele count and total allele number (read depth) across all samples.</u>

#### Quantifying plausible environmental correlates of global w*Mel* frequency variation

We estimated several temperature metrics and other environmental variables that could plausibly influence *w*Mel frequencies. We acquired data from NASA POWER (Sparks 2018), which provides temperature and precipitation measurements aggregated for 1981–2023 at 1800 arc second resolution (∼50 km at the equator). We calculated long-term climatological estimates of the 19 representative bioclimatic environmental variables for each location represented in our dataset. We used long-term climatological means to test the influence of stable environmental gradients on *w*Mel frequency rather than stochastic yearly weather variations. While measurements are unlikely to reflect the specific environments experienced by fly hosts, they provide the opportunity to identify long-term environmental correlates of *w*Mel frequency that may contribute to observed frequency clines (Kriesner et al. 2016). We contrast patterns that we observed for the shorter-term *w*Mel frequency and temperature estimates described above for orchard populations.

### Statistical analyses

All statistical analyses were conducted in R (R Core Team 2020). We estimated weekly *w*Mel prevalence for each outdoor cage population in our experimental orchard, the two natural orchard populations, and all other locations from our global dataset as the proportion of *w*Mel-positive individuals. Exact 95% confidence intervals for *w*Mel frequencies were calculated using the *Hmisc* package (Harrell Jr 2025).

#### Analyzing wMel frequencies in experimental and natural orchard populations

To test for spatiotemporal variation in *w*Mel frequency, we used generalized additive models (GAM) using the *gam()* function with penalized spline on week of sampling to capture the overall temporal trend, a parametric term for cage or location identity, and a tensor product interaction smooth between week and cage or location identity to capture group-specific deviations using a restricted maximum likelihood (REML) method from the *mgcv* package (Wood 2003; Wood 2004; Wood 2011; Wood et al. 2016; Wood 2017). Models were fit using binomial errors with paired *w*Mel-positive and *w*Mel-negative counts as the dependent variable and week of sampling, cage identity, and their interactions as independent variables for the experimental orchard in our first set of regression models. We compared the full model with a null model without cage identity to assess significant spatial variation.

We ran a second set of regression models using paired *w*Mel-positive and *w*Mel-negative counts as the dependent variable with week of sampling, location identity, and their interactions as independent variables for testing spatiotemporal variation in *w*Mel frequency in the two natural orchards. Penalized splines were applied to continuous predictors, with tensor product smooths used for interactions between time and cage or location identity. We assessed basis adequacy by comparing AIC scores (Burnham et al. 2011) and using the *gam.check()* function by fitting models across different k values ranging from 3 to 14 (the number of unique weeks sampled). Smooth estimates of individual model components were extracted using the *gratia ()* package (Simpson 2024).

For temperature data in the experimental orchard, we first tested if weekly mean and weekly average maximum temperature among HOBO loggers in focal experimental cages differed significantly, using Gaussian GAM models with temperature as the dependent variable with week as a continuous independent variable and logger identity as a fixed effect. We compared the full model with a null model without logger identity. Weekly mean and weekly average maximum temperatures did not differ significantly among loggers (weekly mean: GAM LRT = 8.542, *P* = 0.66; weekly average maximum: GAM LRT = 20.78, *P* = 0.43). Hence, we pooled weekly mean and weekly average maximum temperature data across loggers and used the pooled data for downstream analysis of experimental orchard data. To validate the use of NSRDB temperature data for sites lacking on-ground sensors, we compared weekly mean and weekly average maximum temperatures from HOBO loggers deployed at the experimental orchard against corresponding NSRDB estimates for the same location and weeks. HOBO and NSRDB temperatures were strongly correlated for both weekly mean (Pearson r = 0.997, GAM R² = 0.994, *P* < 0.001) and weekly average maximum (Pearson r = 0.990, GAM R² = 0.985, *P* < 0.001), confirming that NSRDB data reliably captured the ambient thermal conditions at our study sites.

To test the influence of temperature on *w*Mel frequency variation at our experimental cages and at the natural orchards, we fit binomial GAMs with paired *w*Mel-positive and *w*Mel-negative counts as the dependent variable. We used scaled temperature (weekly mean and weekly average maximum) lagged by one week (lag-1), corresponding to the temperatures experienced by females during oviposition as independent variable, and cage identity as random effect smoothing (*re*) for the experimental orchard observations. Mean egg-to-eclosion development time across all three locations was 9.9 days (range: 7.3–10.8 days across months), motivating a one-week lag as a close approximation to a single generation. Weekly and 10-day lagged temperatures were strongly correlated for both metrics (Pearson *r* = 0.91 and 0.93, respectively), and GAM predictions were qualitatively identical across both lag windows (**Fig. S8**).

For the experimental cages, a penalized spline was applied to temperature acquired from HOBO loggers. For comparing *w*Mel frequency between the natural orchards, location identity was included as both a parametric term and a tensor product interaction with random effect smoothing (*re*), assuming a shared functional response to temperature estimates from NSRDB across locations while allowing differences in magnitude. This contrasts with the spatiotemporal models above, where factor-smooth interactions (bs = “*fs*”) were used to allow cage- and location-specific temporal trajectories to differ in shape. Similar models were constructed with scaled average maximum temperature lagged by one week (lag-1) with the same set of independent variables. The experimental orchard model was refit with quasi-binomial errors as the binomial model exhibited significant overdispersion (dispersion= 1.15, *P*=0.03). We checked for basis adequacy using the *gam.check()* function. We defined peak thermal ranges as the temperature interval within which the GAM smooth remained within 5% of its maximum predicted value. In addition, we fit a binomial model to test if *w*Mel frequencies varied seasonally and temporally at Linvilla, PA for the long-term observations using the *glm ()* function. We used paired *w*Mel-positive and *w*Mel-negative counts as the dependent variable and used month as independent variable for short-term variation and year of sampling as independent factor for long-term variation.

#### Analyzing global wMel frequency variation

We first replicated the logistic regression framework of Kriesner et al. (2016) using *glm()* on their original dataset and continent groups, with paired *w*Mel-positive and *w*Mel-negative counts as the response and absolute latitude as the predictor. Model diagnostics via *testDispersion()* and *testZeroInflation()* in the DHARMa package (Hartig et al. 2024) revealed significant overdispersion, zero-inflation, and quantile deviations for almost all models (see SI). Their *glm()* framework also precludes random effects, leaving location-specific non-independence unaccounted for. Hence, we used Bayesian beta-binomial regression, which accommodates overdispersion through the beta-binomial likelihood and accounts for non-independence through partial pooling of location and year random intercepts. The temporal component of *w*Mel frequency variation is linked to temperature at the orchard scale, but the global dataset lacks the resolution to model it mechanistically, so the year-within-location intercept absorbs it structurally. Partial pooling also handles locations with small sample sizes by shrinking their frequency estimates toward the global mean, ensuring they contribute proportionally less to fixed-effect inference without biasing it. Our expanded dataset therefore includes locations with fewer than 10 individuals that were appropriately excluded by Kriesner et al. using their framework.

We fit two Bayesian beta-binomial model sets: one testing the relationship between *w*Mel frequency and latitude, and one identifying specific bioclimatic predictors. Because bioclimatic variables are partly collinear with latitude, and combining latitude splines with bioclimatic terms in a single model produces competing predictors that are not cleanly interpretable, we fit these models separately. Both model sets used continent as the geographic grouping variable, consistent with Kriesner et al. (2016), but we defined five continental groups (Africa, Asia, Australia, Europe, and the Americas) rather than their four (Africa, Australia, Eurasia, and North America). South American observations (6 observations from 3 unique latitudes) were not included in their dataset and were merged with North America in ours because the data were too sparse to support a separate continent-level effect. In contrast, ample data exist to split Eurasia into Asia and Europe, which we did based on biological differences relevant to our study—European *D. melanogaster* populations likely persist year-round, whereas central Asian populations may experience seasonal recolonization (Ilinsky and Zakharov 2007; Kriesner et al. 2016) that influences *w*Mel frequency dynamics.

We fit all Bayesian models using the brms package (Bürkner 2017) with four chains of 4000 iterations each (1000 warmup; 12,000 post-warmup draws). We specified weakly informative priors: normal (0, 1) on fixed-effect coefficients, normal (0, 1.5) on the intercept, exponential (2) on all random effect standard deviations, and exponential (2) on the beta-binomial precision parameter (φ). Convergence was confirmed by Rhat < 1.01 for all parameters, adequate bulk and tail effective sample sizes, and the absence of divergent transitions. Model comparison was performed using approximate leave-one-out cross-validation (LOO) via the loo package (Vehtari et al. 2017), with ELPD differences and standard errors reported following (Sivula et al. 2025).

For the latitude analysis, we fit two models. For the Kriesner et al. (2016) dataset, we fit a model with a linear term for scaled absolute latitude and random intercepts for location and year nested within location; the data were too sparse to support continent-specific splines. For our expanded dataset, we fit a model with continent-specific spline terms for scaled absolute latitude, continent as a random intercept, and random intercepts for location and year nested within location. We evaluated spline basis adequacy by fitting models across basis dimensions ranging from 3 to 40 and selecting the value that maximized ELPD.

To identify specific bioclimatic predictors, we tested pairwise correlations among the 19 bioclimatic variables, grouped those with |*r*| > 0.8 into correlated sets, and retained one representative per set based on biological relevance (**Table S17**). For retained variables significantly correlated with absolute latitude, we extracted residuals from a GAM of the variable on absolute latitude to isolate climatic variation independent of geography. We fit a Bayesian beta-binomial model to this reduced set consisting of seven bioclimatic variables using regularized horseshoe priors (Piironen and Vehtari 2017) on all location-level coefficients, with continent, location, and year nested within location as random intercepts. Variables whose 95% credible intervals excluded zero were retained for the bioclimatic model. The horseshoe retained four predictors: residuals of scaled BIO6 (minimum temperature of the coldest month), scaled BIO8 (mean temperature of the wettest quarter), scaled BIO15 (precipitation seasonality), and scaled BIO17 (precipitation of the driest quarter). The four predictors were robust to alternative model specifications, including models that used hemisphere rather than continent as the grouping variable.

We refit the bioclimatic model with standard priors on the four retained predictors. The model included residuals of scaled BIO6, scaled BIO8, scaled BIO15, and scaled BIO17 as global fixed effects, continent as a random intercept, and a continent-specific random slope for BIO8 estimated without intercept-slope correlation. BIO8 measures mean temperature of the wettest quarter, but the wettest quarter falls in different seasons across continents: summer in much of Australia, winter or spring in Europe, and variable timing across North America and Asia. The same BIO8 value therefore maps onto ecologically different conditions depending on continent. This random slope was specified a priori on these biological grounds, not from inspecting model output, and was confirmed by LOO (ΔELPD = 8.1, SE = 4.7). BIO6, BIO15, and BIO17 measure quantities with consistent biological meaning regardless of continent and were modeled with shared global fixed effects. We included random intercepts for location and year nested within location. We compared models with spline versus linear terms for BIO6, BIO15, and BIO17 using LOO; the spline model did not improve predictive performance (|ΔELPD| < 4 Sivula et al. 2025), so linear terms were retained. Posterior predictive checks indicated adequate model fit (**Fig. S9**).

Finally, we quantified temporal *w*Mel frequency changes at locations with repeated sampling (≥3 observations), categorized as short-term (multiple samples within a single year) or long-term (samples spanning multiple years). For each location, we fit binomial GLMs with paired *w*Mel-positive and *w*Mel-negative counts as the response, using month of sampling as a fixed factor for short-term analyses and year of sampling for long-term analyses. Models exhibiting overdispersion were refit with quasi-binomial errors. Hoffmann et al. (1998) assessed temporal variation using G-tests, which test for heterogeneity (*i.e.,* did frequencies change), whereas our GLMs test for systematic temporal patterns. Hence, we also used G-tests to assess frequency change.

#### Analyzing wMel SNP-environment associations and lineage structure

While *w*Mel prevalence could not be estimated from pooled samples, allele frequencies among *w*Mel-positive reads can be estimated directly from read counts at each position. To identify SNPs associated with latitudinal changes and climatic factors, we used a logit-linked beta-binomial generalized linear mixed model implemented in the *glmmTMB ()* package (McGillycuddy et al. 2025). We retained SNPs present in ≥10 populations and polymorphic in ≥3 populations. For each SNP, we tested for association between allele frequency and each of five environmental predictors: scaled absolute latitude, residuals of scaled BIO6, scaled BIO8, scaled BIO15, and scaled BIO17. Each model included hemisphere identity and sequencing strategy (individual or pooled) as fixed effects and population identity as a random intercept. We used hemisphere rather than continent as a geographic covariate because too few populations were sampled from Australia and Africa to support continent-level effects in the SNP models. Significance was assessed using a likelihood ratio test (LRT) comparing the full model to a null model excluding the predictor.

To account for inflated false positives due to extensive linkage disequilibrium across the *w*Mel genome, we performed 10,000 permutations per predictor in which predictor values were randomly reassigned among populations within sequencing strategy and hemisphere, repeating the complete genome scan for each permuted dataset. Genome-wide significance thresholds were defined as the 95th percentile of the permuted maximum-LRT distribution, controlling the family-wise error rate. We also computed an empirical genome-wide *P*-value for each predictor as the proportion of permutations in which the permuted maximum LRT equaled or exceeded the observed maximum. We tested for spatial autocorrelation in *w*Mel allele frequencies using Moran’s I (Paradis et al. 2004), computed on mean allele frequencies per population against an inverse-distance weight matrix. Some sampling localities are geographically proximate, but the permutation scheme accounts for this spatial non-independence because predictor values are reassigned among populations while the allele frequency data structure – including any shared signal among nearby sites – is preserved; population random intercepts provide additional correction.

To test whether latitude-associated SNPs reflected *w*Mel lineage structure rather than independent allele-environment associations, we repeated the genome scan replacing hemisphere with covariates that capture progressively finer scales of lineage structure. We tested an Africa/non-Africa binary covariate and a Europe/non-Europe binary covariate, each included as a fixed effect in the same beta-binomial GLMM framework with significance assessed using 10,000 permutations per covariate. We also computed the first principal component (PC1) of allele frequencies from the 555 SNPs not associated with latitude, which we used as a fixed-effect covariate. To test whether any latitude signal remained within continents, we ran separate scans restricted to European populations (95 populations, ∼35°–60°N) and North American populations (22 populations, ∼25°–47°N), retaining SNPs present in ≥5 populations and polymorphic in ≥ 2 populations (to account for lower sample sizes compared to the global scan) using the same model structure without the geographic covariate.

We estimated linkage disequilibrium (LD) between all pairs of *w*Mel SNPs from the 339 individually sequenced genomes using the statistic *D’* (Lewontin 1964), where *D’* = 0 indicates no LD and |*D’*| = 1 indicates complete LD. Because recombination erodes LD as a function of distance, decay of LD with physical distance is evidence for recombination (Awadalla et al. 1999). We tested for such correlations using |D’|, but also r² (Hill and Robertson 1966), which is more robust to variation in mutation rates (Awadalla et al. 1999; Meunier and Eyre-Walker 2001; Innan and Nordborg 2002). We additionally tested for the presence of all four haplotype combinations at pairs of SNPs (the four-gamete test; Hudson and Kaplan 1985). We partitioned SNP pairs by genomic compartment – core genome versus WO prophage – with prophage boundaries (Kent et al. 2011; LePage et al. 2017).

We reconstructed *w*Mel genealogical relationships using two complementary approaches. We inferred a *w*Mel haplotype network from the 593 SNP positions across all 339 individually sequenced genomes using the minimum spanning network implemented in the pegas package (Paradis 2010) in R. To reconstruct the *w*Mel genealogy, we identified 728 single-copy protein-coding genes of equal length in the *w*Mel reference (clade III; GCF_000008025.1) and the *w*Zts reference (GCF_032849005.1) from *Z. tsacasi* (Shropshire et al. 2026), extracted orthologous sequences from each of the 339 individually sequenced *w*Mel genomes, and concatenated the alignment (736,557 bp). We inferred a maximum-likelihood phylogeny using RAxML (Stamatakis 2014) using the GTR + Γ model partitioned by codon position with 500 bootstrap replicates, rooted on *w*Zts. Cytoplasmic group identity of *w*Mel lineages was assigned based on the classification framework established by Richardson et al. (2012). Representative sequences from their dataset were included in our phylogenetic tree, and clade membership was inferred by the topological position of our sequences relative to these references. For novel sequences that did not fall within any established clade, we used the set of clade-diagnostic SNPs defined by Richardson et al. and compared each unclassified sequence to the diagnostic profiles of all known clades, assigning clade identity based on the minimum number of mismatches.

A group of sequences comprising samples from southwestern Europe formed a distinct clade that was not assigned to clades I–IV and clade VI. Richardson et al. (2012) assigned two uninfected French strains to clade V based solely on host mitochondrial haplotype. We therefore compared the mitochondrial SNPs of our unassigned sequences against the diagnostic profiles of all Richardson et al. clades. These sequences showed the fewest mismatches with clade V, supporting their assignment to this clade. One additional group of sequences could not be resolved using the Richardson et al. framework. Chrostek et al. (2013) described a novel clade, clade VIII (*w*Mel2), comprising sequences from the Far East, specifically Japan and China, characterized by distinct phylogenomic placement and a duplication of the WO-B prophage region (positions 569,001–634,000 of the *w*Mel reference). For these sequences, we quantified mean sequencing read depth across the WO-B region and normalized it against mean depth across a 250,000 bp window in the core genome. The five Beijing sequences showed WO-B-to-core depth ratios of approximately 1.87–1.92, consistent with a single additional copy of the WO-B region and clearly distinct from all other samples (ratios 0.80–1.11), and were accordingly assigned to clade VIII.

We characterized the functional composition of significant and background SNPs in several ways. We used SnpEff (Cingolani et al. 2012) to annotate predicted functional effects, annotating the strongest effect for each SNP (high > moderate > low > modifier), as a single SNP can have multiple functional effects. For SNPs assigned as a modifier, we reported the nearest gene for biological context. We then tested whether latitude-associated SNPs were enriched in the WO prophage region relative to the full SNP set using Fisher’s exact test. We tested whether nonsynonymous sites were enriched among latitude-associated SNPs relative to background using a 2 × 2 Fisher’s exact test and compared the folded site-frequency spectra of nonsynonymous and synonymous sites across all 593 biallelic SNPs using a Kolmogorov-Smirnov test. Per-SNP frequencies were computed as equal-weighted means across populations using individually sequenced genomes, and SNPs classified as intergenic, upstream/downstream, or modifier were excluded from functional-class comparisons.

To test whether the skew in the site-frequency spectrum toward rare alleles differed between nonsynonymous and synonymous sites, we implemented the heterogeneity test of Hahn et al. (2002). From 339 individually sequenced *w*Mel genomes, we retained sites with data from ≥335 individuals and computed Tajima’s *D* separately for each class, subsampling allele counts to *N* = 335 via hypergeometric draws where necessary. We evaluated significance using 10,000 coalescent simulations with no recombination conditioned on observed *S* for each class, implemented in msprime (Baumdicker et al. 2022). The two-tailed *P*-value was the proportion of simulated |Δ*D*| ≥ observed |Δ*D*|. To polarize derived allele frequencies, we aligned the *w*Ri reference to the *w*Mel reference using minimap2 with default parameters (Li 2018), extracting per-position alleles with SAMtools mpileup (Li et al. 2009). SNPs where the outgroup allele matched neither *w*Mel allele were excluded. We compared unfolded site-frequency spectra of nonsynonymous and synonymous sites using the Kolmogorov-Smirnov test.

## Data availability

All data used in this study will be deposited in the Dryad Digital Repository upon acceptance for publication.

## Funding

This work was supported by National Science Foundation (NSF) CAREER (2145195) and National Institutes of Health MIRA (R35GM124701) Awards to BSC. An NIH Award (R01GM137430) supported PRS. Computational resources were provided by the Advanced Cyberinfrastructure Coordination Ecosystem: Services & Support (ACCESS) program through allocation BIO240156 awarded to NR, supported by U.S. National Science Foundation grants #2138259, #2138286, #2138307, #2137603, and #2138296. ELB was supported by an NSF Graduate Research Fellowship Program #DGE-0822 for this work. Additional support was provided by the University of Montana Genomics Core (UMGC) and Montana INBRE Data Science Core, which are funded by the National Institute of General Medical Sciences (P20GM103474), the Office of the Vice President for Research and Creative Scholarship at the University of Montana, and the M. J. Murdock Charitable Trust (202324717 to BSC).

## Notes

### Competing Interest Statement

The authors have declared no competing interest.

