## Supplementary information for "Climate predicts *w*Mel *Wolbachia* frequency variation in *Drosophila melanogaster*, but genomic variation reflects a recent incomplete cytoplasmic sweep"

**Supplemental Material**

**Kriesner et al. (2016) dataset deduplication**

We identified 356 duplicated individual observations (26 spatiotemporal populations, 12 unique locations) in the dataset curated by Kriesner et al. (2016) resulting from overlap among three studies (Verspoor and Haddrill 2011; Ilinsky 2013; Webster et al. 2015), reducing the dataset from 7,469 individuals (173 spatiotemporal populations, 116 unique locations) to 7,113 individuals (147 spatiotemporal populations, 104 unique locations). Duplications were restricted to three locations in Central Asia, four in Africa, and five in Europe. Kriesner et al.’s GLM analyses did not include these continents, but our Bayesian analyses do. All of our analyses related to Kriesner et al. use the deduplicated dataset (*i.e.,* Kdata). Our expanded dataset (Edata) combined Kdata with novel genomic and experimental data collected since their study, resulting in frequency estimates derived from 13,199 individuals and 434 spatiotemporal collections spanning 248 locations across 5 continents and 42 years of sampling (**Fig. 3A**).

**GLM replication of Kriesner et al. (2016)**

We replicated the logistic GLM framework of Kriesner et al. (2016) on Kdata using their sub-analysis criteria. Full GLM results are presented in **Tables S13**. DHARMa diagnostics (testDispersion, testZeroInflation) indicated significant overdispersion and quantile deviations for all models across both datasets (**Table S14**). These diagnostic failures, combined with the absence of random effects in the GLM framework, motivated our use of Bayesian beta-binomial regression for all subsequent analyses.

**Bioclimatic variable selection and model comparison**

We tested pairwise correlations among the 19 WorldClim bioclimatic variables and grouped those with |*r*| > 0.8 into correlated sets, retaining one representative per set based on biological relevance (**Table S17**). For retained variables significantly correlated with absolute latitude, we extracted residuals from a GAM of the variable on absolute latitude to isolate climatic variation independent of geography. Of the variables, BIO1 (annual mean temperature), BIO3 (isothermality), BIO6 (minimum temperature of the coldest month), and BIO11 (mean temperature of the coldest quarter) were each strongly correlated with absolute latitude (Pearson's r = −0.864, −0.833, −0.836, and −0.878, respectively) and were residualized on absolute latitude prior to downstream analysis (GAM R² = 0.86, 0.73, 0.77, and 0.85, respectively).

Our regularized horseshoe model tested seven bioclimatic predictors after collinearity filtering. Four predictors had 95% credible intervals excluding zero and were retained for the bioclimatic model: residuals of scaled BIO6 (est.: −0.44 [−0.78, −0.09]; bulk ESS=7477, tail ESS=6572), scaled BIO8 (est.: 0.21 [0.02, 0.40]; bulk ESS=8173, tail ESS=6255), scaled BIO15 (est.: 0.44 [0.04, 0.80]; bulk ESS=5381, tail=3263), and scaled BIO17 (est.: 0.37 [0.01, 0.69]; bulk ESS=5620, tail ESS=3567). Three predictors were not retained: residuals of scaled BIO3 (est.: 0.10 [−0.09, 0.36]; bulk ESS=6561, tail ESS=10169), scaled BIO10 (est.: 0.01 [−0.14, 0.16]; bulk ESS=8862, tail ESS=10436), and scaled BIO16 (est.: 0.04 [−0.16, 0.27]; bulk ESS=7192, tail ESS=6563). BIO3 (isothermality) was not retained because continent random intercepts absorbed the geographic structure it was capturing.

We compared the bioclimatic model with and without a continent-specific random slope for BIO8. The random slope model was favored (ΔELPD = 8.1, SE = 4.7). We also compared models with linear versus spline terms for BIO6, BIO15, and BIO17; the spline model did not improve predictive performance (|ΔELPD| < 4). BIO8 was retained as a linear term because it already included a continent-specific random slope. ELPD differences and standard errors are reported following Sivula et al. (2025). All Bayesian models converged with Rhat < 1.01 for all parameters, minimum bulk ESS (1482), minimum tail ESS of (2515), and no divergent transitions.

**Temporal *w*Mel frequency stability**

Full temporal stability results for all 28 locations are in **Table S15**. G-tests consistently detected more temporal variation than GLMs, as expected given that G-tests assess any frequency heterogeneity across sampling periods whereas GLMs test for systematic temporal patterns and require larger sample sizes to achieve significance. The majority of locations showed no significant temporal variation by either test, indicating that *w*Mel frequencies are broadly stable through time at most sites. These results are broadly consistent with Kriesner et al. (2016), who reported general temporal stability in Australian populations with notable exceptions at Cobram and Gold Coast, both of which showed significant variation in our analyses. Hastings showed significant variation by GLM alone in our analysis (quasibinomial *P* = 0.018) but not by G-test (*P* = 0.138), contrasting with Hoffmann et al. (1998) who found no significant variation at this location.

**Descriptive temperature–frequency visualization**

Descriptive visualization of raw *w*Mel frequencies against mean temperature of the warmest quarter (BIO10; **Fig. S4**) illustrates the temperature–frequency landscape that the bioclimatic model decomposes into specific predictive axes. BIO10 was not retained as a predictor in the formal bioclimatic model, but captures the active *D. melanogaster* reproductive season more directly than annual mean temperature (BIO1). *w*Mel frequencies increase from ~17°C to ~24°C before declining at higher temperatures, consistent with intermediate temperatures favoring maternal transmission (Hague et al. 2022) and the orchard-scale temperature associations documented above. The large-sample, low-frequency observation at ~27°C corresponds to the Ghana population, for which Kriesner et al. (2016) also noted anomalously low equatorial frequencies.

**Supplementary Tables**

| **Table S1**. Sample sizes and mean *w*Mel *Wolbachia* frequency estimates with exact 95% binomial confidence (in brackets) for each week across all cages sampled in the UPenn experimental orchard. | | | | | |
| --- | --- | --- | --- | --- | --- |
| **Week number** | | ***w*Mel Positive (*N*)** | **Total (*N*)** | | **Mean Frequency [95% CI]** |
| 1 | 170 | | | 202 | 0.84 [0.78-0.89] |
| 2 | 139 | | | 186 | 0.75 [0.68-0.81] |
| 3 | 97 | | | 134 | 0.72 [0.64-0.80] |
| 4 | 154 | | | 177 | 0.87 [0.81-0.92] |
| 5 | 192 | | | 217 | 0.88 [0.83-0.92] |
| 6 | 180 | | | 193 | 0.93 [0.89-0.96] |
| 7 | 124 | | | 145 | 0.86 [0.79-0.91] |
| 8 | 119 | | | 135 | 0.88 [0.81-0.93] |
| 9 | 160 | | | 181 | 0.88 [0.83-0.93] |
| 10 | 153 | | | 201 | 0.76 [0.70-0.82] |
| 11 | 175 | | | 196 | 0.89 [0.84-0.93] |
| 12 | 174 | | | 223 | 0.78 [0.72-0.83] |
| 13 | 119 | | | 164 | 0.73 [0.65-0.79] |
| 14 | 189 | | | 209 | 0.9 [0.86-0.94] |

**Table S2**. Sample sizes and mean *w*Mel *Wolbachia* frequency estimates with exact 95% binomial confidence (in brackets) for each cage across all weeks sampled at the UPenn experimental orchard.

| **Cage number** | ***w*Mel Positive (*N*)** | **Total (*N*)** | **Mean Frequency [95% CI]** |
| --- | --- | --- | --- |
| E01 | 211 | 237 | 0.89 [0.84-0.93] |
| E02 | 206 | 240 | 0.86 [0.81-0.9] |
| E03 | 197 | 227 | 0.87 [0.82-0.91] |
| E04 | 159 | 204 | 0.78 [0.72-0.83] |
| E05 | 167 | 203 | 0.82 [0.76-0.87] |
| E06 | 183 | 225 | 0.81 [0.76-0.86] |
| E07 | 193 | 226 | 0.85 [0.8-0.9] |
| E08 | 175 | 220 | 0.8 [0.74-0.85] |
| E09 | 180 | 216 | 0.83 [0.78-0.88] |
| E10 | 166 | 201 | 0.83 [0.77-0.88] |
| E11 | 160 | 186 | 0.86 [0.8-0.91] |
| E12 | 148 | 178 | 0.83 [0.77-0.88] |

**Table S3**. *w*Mel *Wolbachia* frequency estimates with exact 95% binomial confidence (in brackets) and sample sizes (in parentheses). Weekly frequency estimates are presented for each cage (E01–E12) sampled at the UPenn experimental orchard. Blank cells represent unsampled weeks.

| **Week number** | **E01** | **E02** | **E03** | **E04** | **E05** | **E06** | **E07** | **E08** | **E09** | **E10** | **E11** | **E12** |
| --- | --- | --- | --- | --- | --- | --- | --- | --- | --- | --- | --- | --- |
| 1 | 0.94 [0.71-1]; (17) | 0.94 [0.73-1]; (18) | 0.88 [0.62-0.98]; (16) | 0.76 [0.5-0.93]; (17) | 0.84 [0.6-0.97]; (19) | 0.69 [0.41-0.89]; (16) | 0.82 [0.57-0.96]; (17) | 0.76 [0.5-0.93]; (17) | 0.69 [0.41-0.89]; (16) | 0.8 [0.56-0.94]; (20) | 1 [0.8-1]; (17) | 1 [0.74-1]; (12) |
| 2 | 1 [0.83-1]; (20) | 0.94 [0.73-1]; (18) | 0.75 [0.48-0.93]; (16) | 0.78 [0.52-0.94]; (18) | 0.82 [0.57-0.96]; (17) | 0.83 [0.52-0.98]; (12) | 0.53 [0.28-0.77]; (17) | 0.31 [0.09-0.61]; (13) | 0.69 [0.41-0.89]; (16) | 0.62 [0.32-0.86]; (13) | 0.63 [0.35-0.85]; (16) | 1 [0.69-1]; (10) |
| 3 | 1 [0.78-1]; (15) | 0.87 [0.6-0.98]; (15) | 1 [0.74-1]; (12) | 0.94 [0.71-1]; (17) |  | 0.56 [0.3-0.8]; (16) | 0.54 [0.25-0.81]; (13) | 0.54 [0.25-0.81]; (13) | 0.36 [0.13-0.65]; (14) | 0.68 [0.43-0.87]; (19) |  |  |
| 4 | 1 [0.8-1]; (17) | 0.82 [0.57-0.96]; (17) | 0.83 [0.52-0.98]; (12) | 0.92 [0.62-1]; (12) | 1 [0.77-1]; (14) | 0.87 [0.6-0.98]; (15) | 0.82 [0.48-0.98]; (11) | 1 [0.72-1]; (11) | 0.94 [0.71-1]; (17) | 0.94 [0.7-1]; (16) | 0.65 [0.38-0.86]; (17) | 0.72 [0.47-0.9]; (18) |
| 5 | 0.8 [0.56-0.94]; (20) | 1 [0.82-1]; (19) | 0.88 [0.64-0.99]; (17) | 0.89 [0.65-0.99]; (18) | 0.89 [0.65-0.99]; (18) | 0.85 [0.62-0.97]; (20) | 0.88 [0.64-0.99]; (17) | 0.84 [0.6-0.97]; (19) | 0.94 [0.7-1]; (16) | 1 [0.81-1]; (18) | 0.78 [0.52-0.94]; (18) | 0.88 [0.64-0.99]; (17) |
| 6 | 0.94 [0.71-1]; (17) | 1 [0.78-1]; (15) | 1 [0.79-1]; (16) | 0.93 [0.66-1]; (14) | 0.93 [0.66-1]; (14) | 0.82 [0.57-0.96]; (17) | 0.94 [0.73-1]; (18) | 0.75 [0.48-0.93]; (16) | 1 [0.79-1]; (16) | 1 [0.8-1]; (17) | 0.94 [0.73-1]; (18) | 0.93 [0.68-1]; (15) |
| 7 | 0.94 [0.7-1]; (16) | 0.67 [0.38-0.88]; (15) | 0.79 [0.54-0.94]; (19) | 0.8 [0.52-0.96]; (15) | 0.83 [0.59-0.96]; (18) | 0.95 [0.75-1]; (20) | 1 [0.81-1]; (18) | 0.79 [0.54-0.94]; (19) | 1 [0.48-1]; (5) |  |  |  |
| 8 | 0.83 [0.52-0.98]; (12) | 0.83 [0.52-0.98]; (12) | 0.92 [0.62-1]; (12) | 0.67 [0.09-0.99]; (3) | 0.82 [0.48-0.98]; (11) | 1 [0.63-1]; (8) | 0.9 [0.55-1]; (10) | 0.94 [0.7-1]; (16) | 0.77 [0.46-0.95]; (13) | 0.8 [0.44-0.97]; (10) | 1 [0.78-1]; (15) | 0.92 [0.64-1]; (13) |
| 9 | 0.67 [0.35-0.9]; (12) | 0.8 [0.52-0.96]; (15) | 0.93 [0.68-1]; (15) | 0.76 [0.5-0.93]; (17) | 0.82 [0.48-0.98]; (11) | 0.88 [0.64-0.99]; (17) | 1 [0.79-1]; (16) | 0.94 [0.7-1]; (16) | 1 [0.78-1]; (15) | 0.93 [0.68-1]; (15) | 0.94 [0.73-1]; (18) | 0.86 [0.57-0.98]; (14) |
| 10 | 0.8 [0.56-0.94]; (20) | 0.72 [0.47-0.9]; (18) | 0.63 [0.38-0.84]; (19) | 0.53 [0.28-0.77]; (17) | 0.71 [0.44-0.9]; (17) | 0.71 [0.44-0.9]; (17) | 0.76 [0.5-0.93]; (17) | 0.81 [0.54-0.96]; (16) | 0.93 [0.68-1]; (15) | 0.82 [0.57-0.96]; (17) | 1 [0.75-1]; (13) | 0.8 [0.52-0.96]; (15) |
| 11 | 1 [0.81-1]; (18) | 0.94 [0.73-1]; (18) | 1 [0.81-1]; (18) | 0.64 [0.35-0.87]; (14) | 0.89 [0.65-0.99]; (18) | 0.94 [0.7-1]; (16) | 0.95 [0.74-1]; (19) | 1 [0.82-1]; (19) | 0.84 [0.6-0.97]; (19) | 0.79 [0.49-0.95]; (14) | 1 [0.63-1]; (8) | 0.67 [0.38-0.88]; (15) |
| 12 | 0.78 [0.52-0.94]; (18) | 0.73 [0.52-0.88]; (26) | 0.78 [0.56-0.93]; (23) | 0.78 [0.52-0.94]; (18) | 0.65 [0.38-0.86]; (17) | 0.73 [0.5-0.89]; (22) | 0.91 [0.72-0.99]; (23) | 0.86 [0.42-1]; (7) | 0.83 [0.63-0.95]; (24) | 0.82 [0.48-0.98]; (11) | 0.74 [0.49-0.91]; (19) | 0.8 [0.52-0.96]; (15) |
| 13 | 0.78 [0.52-0.94]; (18) | 0.88 [0.64-0.99]; (17) | 0.93 [0.68-1]; (15) | 0.38 [0.09-0.76]; (8) | 0.73 [0.39-0.94]; (11) | 0.75 [0.43-0.95]; (12) | 0.79 [0.49-0.95]; (14) | 0.58 [0.33-0.8]; (19) | 0.82 [0.48-0.98]; (11) | 0.54 [0.25-0.81]; (13) | 0.88 [0.47-1]; (8) | 0.61 [0.36-0.83]; (18) |
| 14 | 0.94 [0.71-1]; (17) | 0.88 [0.64-0.99]; (17) | 0.94 [0.71-1]; (17) | 0.88 [0.62-0.98]; (16) | 0.78 [0.52-0.94]; (18) | 0.88 [0.64-0.99]; (17) | 1 [0.79-1]; (16) | 0.95 [0.74-1]; (19) | 0.89 [0.67-0.99]; (19) | 0.89 [0.65-0.99]; (18) | 0.89 [0.67-0.99]; (19) | 0.94 [0.7-1]; (16) |

**Table S4**. Summary of a binomial generalized additive model (GAM) with logit link examining *w*Mel prevalence as the dependent variable, fit to data from the UPenn experimental orchard only (see **Table S3** for sample sizes). Cage identity (reference level: E01), week number, and their tensor product interaction were included as model terms to test for cage-level differences in baseline prevalence and cage-specific temporal trajectories across 14 consecutive sampling weeks. Parametric coefficients for each cage represent contrasts in baseline prevalence relative to E01 on the logit scale. The main smooth term s(week number) captures the average temporal trajectory across all cages, and the tensor product interaction ti(week number × cage) tests whether temporal trajectories differ significantly among cages. Estimated degrees of freedom (edf) indicate smooth complexity, with the large edf for the interaction term (24.9) reflecting substantial among-cage variation in temporal dynamics, consistent with the cage-specific patterns shown in Fig. S2.

| **Parametric coefficients:** | | **Estimate** | **Std. Error** | **z value** | **Pr(>\|z\|)** |
| --- | --- | --- | --- | --- | --- |
| (Intercept) | 2.159 | | 0.216 | 10.01 | <0.001 |
| cageE02 | | -0.296 | 0.288 | -1.029 | 0.304 |
| cageE03 | | -0.227 | 0.294 | -0.773 | 0.44 |
| cageE04 | | -0.832 | 0.28 | -2.973 | 0.003 |
| cageE05 | | -0.6 | 0.289 | -2.032 | 0.042 |
| cageE06 | | -0.631 | 0.280 | -2.253 | 0.024 |
| cageE07 | | -0.218 | 0.302 | -0.723 | 0.47 |
| cageE08 | | -0.76 | 0.277 | -2.741 | 0.006 |
| cageE09 | | -0.371 | 0.3 | -1.252 | 0.210 |
| cageE10 | | -0.438 | 0.3 | -1.46 | 0.145 |
| cageE11 | | -0.241 | 0.317 | -0.759 | 0.448 |
| cageE12 | | -0.45 | 0.307 | -1.467 | 0.142 |
| **Approx. significance of smooth terms:** | | **edf** | **Ref.df** | **Chi.sq** | ***P* value** |
| spline (week number) | | 7.941 | 9.1 | 37.44 | <0.001 |
| tensor interaction (week number x cage) | | 24.879 | 47 | 67.96 | <0.001 |

**Table S5**. Sample sizes and mean *w*Mel *Wolbachia* frequency estimates with exact 95% binomial confidence (in brackets) across all weeks for each location sampled.

| **Location** | ***w*Mel Positive (*N*)** | **Total (*N*)** | **mean Frequency [95% CI]** |
| --- | --- | --- | --- |
| Linvilla orchard | 211 | 297 | 0.71 [0.66-0.76] |
| Lohr’s orchard | 276 | 391 | 0.71 [0.66-0.75] |
| U. Penn. orchard | 2145 | 2563 | 0.84 [0.82-0.85] |

**Table S6**. Sample sizes and mean *w*Mel *Wolbachia* frequency estimates with exact 95% binomial confidence (in brackets) for each location across each week sampled at the three orchards. The number of individuals sampled are indicated in parentheses.

| **Week number** | **Linvilla orchard** | **Lohr's orchard** | **U. Penn. orchard** |
| --- | --- | --- | --- |
| 1 |  |  | 0.84 [0.78-0.89]; (202) |
| 2 | 0.52 [0.37-0.68]; (44) | 0.73 [0.54-0.87]; (33) | 0.75 [0.68-0.81]; (186) |
| 3 | 0.62 [0.48-0.75]; (55) | 0.64 [0.5-0.76]; (55) | 0.72 [0.64-0.8]; (134) |
| 4 | 0.78 [0.65-0.88]; (55) | 0.58 [0.44-0.72]; (53) | 0.87 [0.81-0.92]; (177) |
| 5 | 0.96 [0.85-0.99]; (45) | 0.73 [0.59-0.84]; (55) | 0.88 [0.83-0.92]; (217) |
| 6 | 0.63 [0.48-0.77]; (46) | 0.66 [0.52-0.78]; (53) | 0.93 [0.89-0.96]; (193) |
| 7 | 0.75 [0.61-0.86]; (52) | 0.63 [0.46-0.78]; (38) | 0.86 [0.79-0.91]; (145) |
| 8 |  |  | 0.88 [0.81-0.93]; (135) |
| 9 |  |  | 0.88 [0.83-0.93]; (181) |
| 10 |  |  | 0.76 [0.7-0.82]; (201) |
| 11 |  | 0.84 [0.64-0.95]; (25) | 0.89 [0.84-0.93]; (196) |
| 12 |  | 0.81 [0.62-0.94]; (27) | 0.78 [0.72-0.83]; (223) |
| 13 |  | 0.84 [0.66-0.95]; (31) | 0.73 [0.65-0.79]; (164) |
| 14 |  | 0.86 [0.64-0.97]; (21) | 0.9 [0.86-0.94]; (209) |

**Table S7**. Summaries of binomial generalized additive models (GAMs) with logit link examining *w*Mel prevalence as the dependent variable, fit separately for each natural population. Week number was included as a smooth term to test for within-season temporal variation in prevalence at each location. For Linvilla, week number spans weeks 2–7; for Lohr's, sampling covered weeks 2–7 and 11–14. The intercept represents baseline prevalence on the logit scale, and estimated degrees of freedom (edf) indicate the complexity of the temporal smooth, where values substantially above 1 indicate non-linearity. Chi-square statistics and associated *P* values test whether week number explains significant variation in prevalence at each location.

| Linvilla |  |  |  |  |
| --- | --- | --- | --- | --- |
| **Parametric coefficients:** | **Estimate** | **Std. Error** | **z value** | **Pr(>\|z\|)** |
| (Intercept) | 0.92 | 0.14 | 7.04 | <0.001 |
| **Approx. significance of smooth terms:** | **edf** | **Ref.df** | **Chi.sq** | ***P* value** |
| spline (week number) | 4.16 | 4.62 | 18.51 | 0.001 |
| Lohr’s |  |  |  |  |
| **Parametric coefficients:** | **Estimate** | **Std. Error** | **z value** | **Pr(>\|z\|)** |
| (Intercept) | 1.02 | 0.13 | 8.09 | <0.001 |
| **Approx. significance of smooth terms:** | **edf** | **Ref.df** | **Chi.sq** | ***P* value** |
| spline (week number) | 1.57 | 1.91 | 9.10 | 0.007 |

**Table S8**. Summary of a binomial generalized linear model (GLM) with logit link examining *w*Mel prevalence as the dependent variable, fit to data from Linvilla orchard, Pennsylvania. Month (reference level: June) and year (reference level: 2009) were included as independent variables to test for seasonal and interannual variation in prevalence. The negative coefficient for month indicates lower prevalence in November (Fall) relative to June (Spring), and year coefficients represent contrasts relative to 2009.

| **Term** | **Estimate** | **Std. Error** | **z value** | **Pr(>\|z\|)** |
| --- | --- | --- | --- | --- |
| Intercept | 0.402 | 0.244 | 1.646 | 0.1 |
| month | -0.552 | 0.176 | -3.144 | 0.002 |
| 2010 | -0.645 | 0.317 | -2.034 | 0.042 |
| 2011 | 0.235 | 0.323 | 0.727 | 0.468 |
| 2012 | 0.077 | 0.323 | 0.238 | 0.812 |
| 2013 | -0.049 | 0.332 | -0.148 | 0.883 |
| 2014 | 0.281 | 0.324 | 0.866 | 0.386 |
| 2015 | 0.037 | 0.334 | 0.111 | 0.911 |

**Table S9**. Summary of quasi-binomial generalized additive models (GAMs) with logit link examining *w*Mel prevalence as the dependent variable, fit to data from the UPenn experimental orchard only (see Table S3 for sample sizes). A quasi-binomial family was used in place of binomial to account for overdispersion, with F-statistics reported accordingly. Models **(a)** and **(b)** differ only in the temperature metric used: lagged weekly mean temperature and lagged weekly average maximum temperature respectively. In both models, the parametric intercept represents the baseline prevalence on the logit scale. The smooth term s(cage) was included to account for cage-level variation but was penalised to zero estimated degrees of freedom (edf = 0) in both models, indicating no residual cage effect after accounting for temperature consistent with the temporal trajectories shown in Fig. S3. Temperature values are lagged by one week to account for *Drosophila* developmental timelines.

| a) lagged weekly (lag-1) mean temperature |  |  |  |  |
| --- | --- | --- | --- | --- |
| **Parametric coefficients:** | **Estimate** | **Std. Error** | **t value** | **Pr(>\|z\|)** |
| (Intercept) | 1.68 | 0.08 | 21.91 | <0.001 |
| **Approx. significance of smooth terms:** | **edf** | **Ref.df** | **F** | ***P* value** |
| spline (lagged weekly mean temperature) | 2.60 | 2.88 | 11.05 | <0.001 |
| spline (cage) | 0 | 11 | <0.001 | 0.57 |
| b) lagged weekly (lag-1) average maximum temperature |  |  |  |  |
| **Parametric coefficients:** | **Estimate** | **Std. Error** | **t value** | **Pr(>\|z\|)** |
| (Intercept) | 1.67 | 0.08 | 21.55 | <0.001 |
| **Approx. significance of smooth terms:** | **edf** | **Ref.df** | **F** | ***P* value** |
| spline (lagged weekly average maximum temp) | 2.63 | 2.90 | 9.79 | <0.001 |
| spline (cage) | 0 | 11 | <0.001 | 0.63 |

**Table S10**. Summary of a binomial generalized additive model (GAM) with logit link examining *w*Mel

prevalence as the response variable, fit to data from all experimental cage and natural populations (see Table S6 for sample sizes). Parametric coefficients represent the intercept (reference level: Linvilla) and contrasts for Lohr's and UPenn relative to Linvilla. Smooth terms include the main effect of lagged weekly mean temperature (lag-1) and its tensor product interaction with location (ti), allowing the magnitude of the temperature response to vary by location. Estimated degrees of freedom (edf) indicate the complexity of each smooth, where values substantially above 1 indicate non-linearity. Chi-square statistics and associated *P* values test whether each smooth term explains significant variation in prevalence. Temperature values are lagged by one week to account for *Drosophila* developmental timelines.

| **Parametric coefficients** | **Estimate** | **Std. Error** | **z value** | **Pr(>\|z\|)** |
| --- | --- | --- | --- | --- |
| (Intercept) | 1.17 | 0.25 | 4.67 | <0.001 |
| Lohr’s | -0.24 | 0.28 | -0.85 | 0.39 |
| UPenn | 0.51 | 0.26 | 1.92 | 0.05 |
| **Approx. significance of smooth terms:** | **edf** | **Ref.df** | **Chi.sq** | ***P* value** |
| spline (lagged weekly mean temperature) | 1.98 | 2 | 59.88 | <0.001 |
| tensor interaction (lagged weekly mean temperature x location) | 1.37 | 2 | 17.48 | <0.001 |

**Table S11**. Summary of a binomial generalized additive model (GAM) with logit link examining *w*Mel prevalence as the response variable, fit to data from all experimental cage and natural populations (see Table S6 for sample sizes). Parametric coefficients represent the intercept (reference level: Linvilla) and contrasts for Lohr's and UPenn relative to Linvilla. Smooth terms include the main effect of lagged weekly average maximum temperature (lag-1) and its tensor product interaction with location (ti), allowing the magnitude of the temperature response to vary by location. Estimated degrees of freedom (edf) indicate the complexity of each smooth, where values substantially above 1 indicate non-linearity. Chi-square statistics and associated *P* values test whether each smooth term explains significant variation in prevalence. Temperature values are lagged by one week to account for *Drosophila* developmental timelines.

| **Parametric coefficients:** | **Estimate** | **Std. Error** | **z value** | **Pr(>\|z\|)** |
| --- | --- | --- | --- | --- |
| (Intercept) | 0.89 | 0.2 | 4.5 | <0.001 |
| Lohr’s | -0.03 | 0.23 | -0.13 | 0.9 |
| UPenn | 0.84 | 0.21 | 3.99 | <0.001 |
| **Approx. significance of smooth terms:** | **edf** | **Ref.df** | **Chi.sq** | **p-value** |
| spline (lagged weekly average maximum temp) | 2.52 | 2.79 | 51.65 | <0.001 |
| tensor interaction (lagged weekly average maximum temp x location) | 1.32 | 2 | 16.26 | <0.001 |

**Table S12**. Weekly mean temperature, weekly average maximum temperature, and *w*Mel frequency across three cage experiment locations (Linvilla, Lohrs, and UPenn). Temperature data for Linvilla and Lohrs were obtained from the NSRDB weather station; temperature data for UPenn were obtained from HOBO data loggers deployed at the cage site.

| **Week number** | **Location** | **Temp mean (°C)** | **Temp max (°C)** | **Mean *w*Mel frequency** |
| --- | --- | --- | --- | --- |
| Week 01 | Linvilla | 25.38 | 30.2 |  |
| Week 02 | Linvilla | 24.97 | 29.56 | 0.52 |
| Week 03 | Linvilla | 25.83 | 31.3 | 0.62 |
| Week 04 | Linvilla | 23.13 | 28.7 | 0.78 |
| Week 05 | Linvilla | 24.6 | 29.64 | 0.96 |
| Week 06 | Linvilla | 24.96 | 30.4 | 0.63 |
| Week 07 | Linvilla | 23.91 | 30.1 | 0.75 |
| Week 08 | Linvilla | 23.05 | 29.33 |  |
| Week 09 | Linvilla | 27.62 | 34.6 |  |
| Week 10 | Linvilla | 21.66 | 27.79 |  |
| Week 11 | Linvilla | 18.37 | 23.89 |  |
| Week 12 | Linvilla | 16.64 | 20.49 |  |
| Week 13 | Linvilla | 19.63 | 25.66 |  |
| Week 14 | Linvilla | 12.83 | 18.64 |  |
| Week 01 | Lohrs | 25.67 | 30.61 |  |
| Week 02 | Lohrs | 25.33 | 29.51 | 0.73 |
| Week 03 | Lohrs | 26.24 | 31.59 | 0.64 |
| Week 04 | Lohrs | 23.5 | 28.9 | 0.58 |
| Week 05 | Lohrs | 24.93 | 29.81 | 0.73 |
| Week 06 | Lohrs | 25.22 | 30.46 | 0.66 |
| Week 07 | Lohrs | 24.63 | 30.61 | 0.63 |
| Week 08 | Lohrs | 23.26 | 29.29 |  |
| Week 09 | Lohrs | 27.66 | 34.23 |  |
| Week 10 | Lohrs | 21.84 | 27.59 |  |
| Week 11 | Lohrs | 18.57 | 23.99 | 0.84 |
| Week 12 | Lohrs | 16.66 | 20.79 | 0.81 |
| Week 13 | Lohrs | 19.35 | 24.99 | 0.84 |
| Week 14 | Lohrs | 13.19 | 18.8 | 0.86 |
| Week 01 | UPenn | 27.61 | 32.6 | 0.84 |
| Week 02 | UPenn | 25.56 | 30.07 | 0.75 |
| Week 03 | UPenn | 26.48 | 30.66 | 0.72 |
| Week 04 | UPenn | 23.39 | 28.73 | 0.87 |
| Week 05 | UPenn | 24.56 | 29.6 | 0.88 |
| Week 06 | UPenn | 25.04 | 29.53 | 0.93 |
| Week 07 | UPenn | 24.14 | 28.87 | 0.86 |
| Week 08 | UPenn | 22.87 | 27.67 | 0.88 |
| Week 09 | UPenn | 27.23 | 33.16 | 0.88 |
| Week 10 | UPenn | 21.38 | 25.63 | 0.76 |
| Week 11 | UPenn | 17.87 | 22.83 | 0.89 |
| Week 12 | UPenn | 16.58 | 19.05 | 0.78 |
| Week 13 | UPenn | 20.03 | 24.86 | 0.73 |
| Week 14 | UPenn | 12.94 | 18.58 | 0.9 |

**Table S13**. **Associations between *w*Mel frequency and latitude across sub-analyses comparing frequentist and Bayesian model frameworks.** Each row represents a geographic sub-analysis replicating Kriesner et al (2016) deduplicated dataset and approach. GLM slope and *P*-value are from binomial generalized linear models (GLMs) with *w*Mel status (positive/negative) as the response and latitude as the predictor analyses includes all observations passing the minimum sample size filter (≥10 individuals screened per location); Bayesian estimates and 95% credible intervals are from beta-binomial mixed models with location and location:year random intercepts, using weakly informative priors (Intercept: Normal(0, 2); slope: Normal(0, 1); random effects SD and overdispersion φ: Exponential(1)). The tropical sub-analysis includes only locations north of −23.5° (within the tropics) in Australia; the temperate sub-analysis includes only locations at or south of −23.5° in Australia. The North America <38°N sub-analysis excludes locations above 38°N latitude. Consistent negative slopes in both Australian sub-analyses across all four model/dataset combinations indicate a robust latitudinal cline in *w*Mel frequency in Australia. The absence of a consistent latitudinal signal in North America across Bayesian models and the GLM suggests the positive slope in the original North American GLM is not robust.

| **Sub-analysis** |  |  | **GLM slope** | **GLM P** |  |  | **Bayesian estimate** | **Bayesian** |
| --- | --- | --- | --- | --- | --- | --- | --- | --- |
| Australia (overall) |  |  | -0.185 | 0 |  |  | -0.119 | [-0.149, -0.089] |
| Australia (tropical, >-23.5°) |  |  | 0.017 | 0.6562 |  |  | 0.031 | [-0.145, 0.213] |
| Australia (temperate, ≤-23.5°) |  |  | -0.227 | 0 |  |  | -0.157 | [-0.21, -0.106] |
| North America (all east coast) |  |  | 0.007 | 0.699 |  |  | -0.013 | [-0.136, 0.106] |
| North America (<38°N) |  |  | -0.221 | 1.00E-04 |  |  | -0.13 | [-0.502, 0.236] |

| **Table S14**. Residual diagnostics for binomial GLMs replicating the analytical approaches of Kriesner et al. (2016), evaluated separately for Australian and regional subsets. "Kdata" refers to the Kriesner et al.’s deduplicated dataset. Dispersion estimates near 1.0 indicate adequate fit; values substantially above 1.0 indicate overdispersion, with the dispersion *P* value testing whether this deviation is significant. The Kolmogorov-Smirnov (KS) statistic and associated *P* value test the uniformity of DHARMa scaled residuals. Quantile deviation *P* values test whether residuals at the 25th (p25), 50th (p50), and 75th (p75) percentiles deviate from expectation, with the combined quantile *P* value providing an overall test across all three. Significant values across dispersion and quantile diagnostics indicate that the binomial GLM approach produces overdispersed and poorly fitting residuals for both datasets. | | | | | | | | |
| --- | --- | --- | --- | --- | --- | --- | --- | --- |
| **Model** | **Dispersion** | **Dispersion *P* value** | **KS stat** | **KS *P* value** | **Quantile p25** | **Quantile p50** | **Quantile p75** | **Quantile combined** |
| Kdata Australia | 1.312 | 0 | 0.254 | 0 | 0.003 | 0 | 0 | 0 |
| Kdata Tropical | 0.976 | 0.684 | 0.205 | 0.153 | 0.708 | 0.752 | 0.06 | 0.179 |
| Kdata Subtropical | 1.214 | 0 | 0.303 | 0 | 0.57 | 0.001 | 0 | 0 |
| Kdata East US | 0.974 | 0.724 | 0.312 | 0.105 | 0.197 | 0.135 | 0.006 | 0.017 |
| Kdata US <38N | 1.037 | 0.618 | 0.405 | 0.107 | 0.657 | 0.213 | 0.065 | 0.196 |

**Table S15**. Results of binomial generalized linear models testing for short-term and long-term temporal variation in *w*Mel frequency at locations with three or more repeated observations. Short-term analyses used month of sampling as the predictor; long-term analyses used year of sampling. Models were fit using a beta-binomial or quasibinomial family depending on overdispersion.

| **Location** | **Temporal scale** | ***N*** | **model** | **GLM *P* value** | **GLM significance** | **G test stat** | **G test *P* value** | **Significance G test** | ***w*Mel frequency variation** |
| --- | --- | --- | --- | --- | --- | --- | --- | --- | --- |
| Cobram, Australia | long_term | 7 | beta-binomial | 0.026 | YES | 21.71 | < 0.001 | YES | SIGNIFICANT (both tests) |
| Alexandrov, Russia | long_term | 4 | quasibinomial | NA (perfect fit) | untestable | 15.27 | 0.002 | YES | G-test only |
| Almaty, Kazakhstan | long_term | 3 | quasibinomial | NA (perfect fit) | untestable | 3.82 | 0.148 | no | not significant |
| Biisk, Altai, Russia | long_term | 3 | quasibinomial | NA (perfect fit) | untestable | 3.11 | 0.212 | no | not significant |
| Bishkek, Kyrgyzstan | long_term | 3 | quasibinomial | NA (perfect fit) | untestable | 0.77 | 0.679 | no | not significant |
| Brest, Belarus | long_term | 3 | quasibinomial | NA (perfect fit) | untestable | 1.72 | 0.424 | no | not significant |
| Cairns, Australia | long_term | 4 | quasibinomial | NA (perfect fit) | untestable | 14.81 | 0.002 | YES | G-test only |
| Dushanbe, Tajikistan | long_term | 3 | quasibinomial | NA (perfect fit) | untestable | 2.88 | 0.237 | no | not significant |
| Krasnodar, Russia | long_term | 5 | quasibinomial | NA (perfect fit) | untestable | 4.87 | 0.301 | no | not significant |
| Kyiv, Ukraine | long_term | 3 | quasibinomial | NA (perfect fit) | untestable | 0.57 | 0.751 | no | not significant |
| Nalchik, Russia | long_term | 7 | quasibinomial | NA (perfect fit) | untestable | 6.86 | 0.334 | no | not significant |
| Tashkent, Uzbekistan | long_term | 6 | quasibinomial | NA (perfect fit) | untestable | 8.31 | 0.14 | no | not significant |
| Linvilla | long_term (also short_term) | 20 | beta-binomial | 0.07 | no | 41.39 | < 0.001 | YES | G-test only |
| Gold Coast, Australia | long_term (also short_term) | 16 | beta-binomial | 0.21 | no | 148.06 | < 0.001 | YES | G-test only |
| Uman, Ukraine | long_term (also short_term) | 18 | quasibinomial | 0.273 | no | 56.03 | < 0.001 | YES | G-test only |
| Coffs Harbour, Australia | long_term (also short_term) | 9 | beta-binomial | 0.569 | no | 41.22 | < 0.001 | YES | G-test only |
| Hastings, Australia | long_term (also short_term) | 5 | quasibinomial | 0.018 | YES | 3.96 | 0.138 | no | GLM/BB only |
| Chernobyl Nuclear Plant, Ukraine | long_term (also short_term) | 4 | quasibinomial | < 0.001 | YES | 21.32 | < 0.001 | YES | SIGNIFICANT (both tests) |
| Odesa, Ukraine | long_term (also short_term) | 8 | beta-binomial | 0.183 | no | 5.77 | 0.123 | no | not significant |
| Brisbane, Australia | long_term (also short_term) | 4 | quasibinomial | 0.391 | no | 1.6 | 0.45 | no | not significant |
| Bowen, Australia | long_term (also short_term) | 3 | quasibinomial | NA (perfect fit) | untestable | 7.89 | 0.019 | YES | G-test only |
| Crete, Greece | long_term (also short_term) | 3 | quasibinomial | NA (perfect fit) | untestable | 36.71 | < 0.001 | YES | G-test only |
| Mackay, Australia | long_term (also short_term) | 3 | quasibinomial | NA (perfect fit) | untestable | 3.18 | 0.204 | no | not significant |
| Nowra, Australia | long_term (also short_term) | 3 | quasibinomial | NA (perfect fit) | untestable | 5.46 | 0.065 | no | not significant |
| Salvador, Bahia, Brazil | long_term (also short_term) | 3 | quasibinomial | NA (perfect fit) | untestable | 2.26 | 0.324 | no | not significant |
| Yarra Valley, Australia | long_term (also short_term) | 3 | quasibinomial | NA (perfect fit) | untestable | 0.91 | 0.634 | no | not significant |
| Yeppoon, Australia | long_term (also short_term) | 3 | quasibinomial | NA (perfect fit) | untestable | 2.17 | 0.339 | no | not significant |
| Lohrs | short_term | 10 | quasibinomial | 0.044 | YES | 12.25 | 0.007 | YES | SIGNIFICANT (both tests) |
| Chernobyl Nuclear Plant, Ukraine | short_term (also long_term) | 4 | beta-binomial | 0.226 | no | 6.69 | 0.01 | YES | G-test only |
| Coffs Harbour, Australia | short_term (also long_term) | 9 | beta-binomial | 0.616 | no | 24.15 | < 0.001 | YES | G-test only |
| Linvilla | short_term (also long_term) | 20 | beta-binomial | 0.005 | YES | 40.33 | < 0.001 | YES | SIGNIFICANT (both tests) |
| Gold Coast, Australia | short_term (also long_term) | 16 | beta-binomial | 0.024 | YES | 269.16 | < 0.001 | YES | SIGNIFICANT (both tests) |
| Uman, Ukraine | short_term (also long_term) | 6 | beta-binomial | 0.048 | YES | 22.56 | < 0.001 | YES | SIGNIFICANT (both tests) |
| Nowra, Australia | short_term (also long_term) | 3 | beta-binomial | 0.17 | no | 2.3 | 0.129 | no | not significant |
| Mackay, Australia | short_term (also long_term) | 3 | quasibinomial | 0.376 | no | 2.21 | 0.137 | no | not significant |
| Bowen, Australia | short_term (also long_term) | 3 | beta-binomial | 0.423 | no | 0.86 | 0.354 | no | not significant |
| Salvador, Bahia, Brazil | short_term (also long_term) | 3 | quasibinomial | 0.442 | no | 1 | 0.318 | no | not significant |
| Yarra Valley, Australia | short_term (also long_term) | 3 | quasibinomial | 0.982 | no | 0 | 0.979 | no | not significant |
| Brisbane, Australia | short_term (also long_term) | 4 | quasibinomial | NA (perfect fit) | untestable | 1.89 | 0.595 | no | not significant |
| Crete, Greece | short_term (also long_term) | 3 | quasibinomial | NA (perfect fit) | untestable | 36.71 | < 0.001 | YES | G-test only |
| Hastings, Australia | short_term (also long_term) | 5 | quasibinomial | NA (perfect fit) | untestable | 4.04 | 0.401 | no | not significant |
| Odesa, Ukraine | short_term (also long_term) | 6 | quasibinomial | NA (perfect fit) | untestable | 15.25 | 0.009 | YES | G-test only |
| Yeppoon, Australia | short_term (also long_term) | 3 | quasibinomial | NA (perfect fit) | untestable | 2.17 | 0.339 | no | not significant |

**Table S16**. Significant SNPs identified by genomic scans based on thresholds derived from 10,000 permuted datasets. SNP effects annotated using SnpEff. Each SNP may generate multiple functional effects depending on the number of genes within the SnpEff annotation window; only the highest-impact effect is reported here (HIGH > MODERATE > LOW > MODIFIER). For MODIFIER effects, the most proximal upstream or downstream gene is reported. Positions indicated by asterisk (*) denote SNPs in the *w*Mel prophage region.

| **Position** | **beta** | **se** | **z** | **LRT stat** | ***P* value LRT** | ***N(*pops)** | ***N* pop (alternate allele present)** | ***N* pop (alternate allele polymorphic)** | ***N* (observed)** | **variable** | **SNP effect** | **Gene name** |
| --- | --- | --- | --- | --- | --- | --- | --- | --- | --- | --- | --- | --- |
| 1066796 | 3.24 | 0.58 | 5.62 | 24.45 | <0.001 | 119 | 9 | 9 | 174 | BIO8 | synonymous variant | ruvB |
| 43974 | 2.14 | 0.5 | 4.26 | 28.88 | <0.001 | 107 | 43 | 42 | 144 | latitude | missense variant | dnaJ |
| 132343 | 1.62 | 0.36 | 4.5 | 30.36 | <0.001 | 100 | 49 | 48 | 128 | latitude | missense variant | gatB |
| 266252 | 2.1 | 0.48 | 4.42 | 27.98 | <0.001 | 82 | 36 | 35 | 93 | latitude | missense variant | ankyrin repeat domain-containing protein |
| 270910 | 1.68 | 0.39 | 4.33 | 30.84 | <0.001 | 97 | 47 | 46 | 125 | latitude | missense variant | ankyrin repeat domain-containing protein |
| 315603 | 1.77 | 0.39 | 4.56 | 34.13 | <0.001 | 103 | 48 | 48 | 130 | latitude | missense variant | Maf family nucleotide pyrophosphatase |
| 357857 | 1.95 | 0.47 | 4.2 | 28.97 | <0.001 | 107 | 38 | 37 | 144 | latitude | missense variant | hypothetical protein |
| 419061 | 2 | 0.45 | 4.47 | 30.54 | <0.001 | 111 | 48 | 48 | 155 | latitude | missense variant | sdhA |
| 493623 | 1.81 | 0.43 | 4.17 | 27.51 | <0.001 | 88 | 38 | 38 | 104 | latitude | missense variant | Rpn family recombination-promoting nuclease/putative transposase |
| 517486 | 1.83 | 0.45 | 4.12 | 25.96 | <0.001 | 92 | 40 | 39 | 117 | latitude | missense variant | pyrH |
| 662811 | 2.13 | 0.49 | 4.31 | 29.58 | <0.001 | 84 | 35 | 34 | 98 | latitude | missense variant | hypothetical protein |
| 805011 | 1.64 | 0.33 | 5.02 | 36.13 | <0.001 | 100 | 53 | 52 | 129 | latitude | missense variant | hypothetical protein |
| 806911 | 2.18 | 0.53 | 4.07 | 28.96 | <0.001 | 113 | 44 | 43 | 152 | latitude | missense variant | uvrB |
| 881800 | 2.14 | 0.51 | 4.19 | 27.83 | <0.001 | 106 | 44 | 44 | 143 | latitude | missense variant | hypothetical protein |
| 1005207 | 1.69 | 0.38 | 4.43 | 29.46 | <0.001 | 104 | 53 | 52 | 131 | latitude | missense variant | hypothetical protein |
| 1054640 | 1.92 | 0.43 | 4.47 | 33.93 | <0.001 | 101 | 48 | 47 | 131 | latitude | missense variant | der |
| 1248515 | 2.21 | 0.49 | 4.5 | 33.72 | <0.001 | 107 | 39 | 38 | 138 | latitude | missense variant | 2-oxoglutarate dehydrogenase E1 component |
| 1253353 | 2.2 | 0.53 | 4.17 | 28.47 | <0.001 | 105 | 45 | 44 | 140 | latitude | missense variant | DsbA family protein |
| 201745* | 1.83 | 0.37 | 4.88 | 36.74 | <0.001 | 103 | 49 | 48 | 135 | latitude | missense variant | PleD family two-component system response regulator |
| 452488* | 2.05 | 0.51 | 4 | 25.29 | <0.001 | 101 | 41 | 40 | 139 | latitude | missense variant | MFS transporter |
| 130710 | 1.81 | 0.36 | 5.02 | 39.43 | <0.001 | 102 | 49 | 48 | 128 | latitude | synonymous variant | hypothetical protein |
| 301562 | 1.9 | 0.43 | 4.37 | 30.42 | <0.001 | 92 | 41 | 40 | 108 | latitude | synonymous variant | tig |
| 717692 | 2.15 | 0.55 | 3.89 | 23.37 | <0.001 | 95 | 35 | 35 | 119 | latitude | synonymous variant | era |
| 929167 | 1.78 | 0.38 | 4.73 | 33.91 | <0.001 | 104 | 49 | 48 | 133 | latitude | synonymous variant | complex I subunit 4 family protein |
| 1154396 | 1.79 | 0.36 | 4.95 | 37.26 | <0.001 | 105 | 48 | 48 | 132 | latitude | synonymous variant | biotin transporter BioY |
| 237268* | 2.15 | 0.47 | 4.61 | 36.21 | <0.001 | 95 | 41 | 40 | 117 | latitude | synonymous variant/ modifier | rmuC/ WO male-killing family protein *Wmk* |
| 163082 | 2.27 | 0.6 | 3.76 | 24.97 | <0.001 | 99 | 39 | 38 | 128 | latitude | upstream/downstream variant | NADH dehydrogenase ubiquinone Fe-S protein 4 |
| 493773 | 2.17 | 0.45 | 4.87 | 31.06 | <0.001 | 88 | 41 | 41 | 108 | latitude | upstream/downstream variant | DNA repair protein pseudogene |
| 585252 | 1.95 | 0.41 | 4.7 | 33.42 | <0.001 | 96 | 44 | 43 | 119 | latitude | upstream/downstream variant | S49 family peptidase |
| 592639 | 1.75 | 0.38 | 4.65 | 32.12 | <0.001 | 97 | 46 | 45 | 120 | latitude | upstream/downstream variant | AAA family ATPase |
| 659208 | 2.12 | 0.53 | 3.97 | 26.02 | <0.001 | 106 | 43 | 43 | 140 | latitude | upstream/downstream variant | rpsC |
| 675549 | 2.08 | 0.52 | 4 | 27.24 | <0.001 | 109 | 44 | 44 | 143 | latitude | upstream/downstream variant | phosphomannomutase |
| 688376 | 2.04 | 0.46 | 4.47 | 35.52 | <0.001 | 95 | 44 | 43 | 114 | latitude | upstream/downstream variant | rpe |
| 812321 | 0.97 | 0.23 | 4.19 | 22.35 | <0.001 | 108 | 65 | 64 | 151 | latitude | upstream/downstream variant | uvrB |
| 1023898 | 1.95 | 0.47 | 4.11 | 26.76 | <0.001 | 105 | 44 | 43 | 143 | latitude | upstream/downstream variant | glutathione S-transferase family protein |
| 1091844 | 2.14 | 0.49 | 4.33 | 25.73 | <0.001 | 107 | 39 | 38 | 143 | latitude | upstream/downstream variant | putative transposase |
| 1181178 | 1.94 | 0.39 | 5 | 38.81 | <0.001 | 97 | 43 | 42 | 121 | latitude | upstream/downstream variant | cold-shock protein |
| 1206452 | 1.78 | 0.36 | 5.01 | 40.67 | <0.001 | 99 | 51 | 49 | 125 | latitude | upstream/downstream variant | OmpA family protein |
| 438970* | 2.5 | 0.63 | 3.96 | 34.43 | <0.001 | 101 | 34 | 33 | 129 | latitude | upstream/downstream variant | phage major capsid protein |

**Table S17**. Pairwise Pearson correlations among 19 scaled WorldClim bioclimatic variables across all locations in the expanded dataset. Variable definitions are given in the footnote. Variables with |r| > 0.8 were grouped into correlated sets and one representative retained per group based on biological relevance. † indicates variables retained after collinearity filtering (*N* = 7); * indicates variables subsequently retained by regularized horseshoe prior variable selection (*N* = 4). Cells with |r| > 0.8 indicate pairs that were grouped during filtering.

|  | **BIO1z** | **BIO2z** | **BIO3z^†^** | **BIO4z** | **BIO5z** | **BIO6z^†*^** | **BIO7z** | **BIO8z^†*^** | **BIO9z** | **BIO10z^†^** | **BIO11z** | **BIO12z** | **BIO13z** | **BIO14z** | **BIO15z^†*^** | **BIO16z^†^** | **BIO17z^†*^** | **BIO18z** | **BIO19z** |
| --- | --- | --- | --- | --- | --- | --- | --- | --- | --- | --- | --- | --- | --- | --- | --- | --- | --- | --- | --- |
| BIO1z | 1 |  |  |  |  |  |  |  |  |  |  |  |  |  |  |  |  |  |  |
| BIO2z | -0.29 | 1 |  |  |  |  |  |  |  |  |  |  |  |  |  |  |  |  |  |
| BIO3z**^†^** | 0.61 | 0.16 | 1 |  |  |  |  |  |  |  |  |  |  |  |  |  |  |  |  |
| BIO4z | -0.84 | 0.41 | -0.75 | 1 |  |  |  |  |  |  |  |  |  |  |  |  |  |  |  |
| BIO5z | 0.57 | 0.37 | 0.21 | -0.09 | 1 |  |  |  |  |  |  |  |  |  |  |  |  |  |  |
| BIO6z**^†*^** | 0.95 | -0.46 | 0.65 | -0.95 | 0.33 | 1 |  |  |  |  |  |  |  |  |  |  |  |  |  |
| BIO7z | -0.8 | 0.62 | -0.6 | 0.97 | 0.02 | -0.94 | 1 |  |  |  |  |  |  |  |  |  |  |  |  |
| BIO8z**^†*^** | 0.48 | -0.28 | 0.15 | -0.26 | 0.32 | 0.4 | -0.31 | 1 |  |  |  |  |  |  |  |  |  |  |  |
| BIO9z | 0.84 | -0.18 | 0.58 | -0.77 | 0.47 | 0.83 | -0.71 | 0 | 1 |  |  |  |  |  |  |  |  |  |  |
| BIO10z**^†^** | 0.79 | -0.05 | 0.21 | -0.32 | 0.9 | 0.59 | -0.3 | 0.51 | 0.6 | 1 |  |  |  |  |  |  |  |  |  |
| BIO11z | 0.97 | -0.36 | 0.69 | -0.94 | 0.39 | 0.99 | -0.9 | 0.4 | 0.85 | 0.62 | 1 |  |  |  |  |  |  |  |  |
| BIO12z | 0.49 | -0.34 | 0.4 | -0.51 | 0.06 | 0.52 | -0.53 | 0.37 | 0.31 | 0.26 | 0.51 | 1 |  |  |  |  |  |  |  |
| BIO13z | 0.56 | -0.29 | 0.43 | -0.54 | 0.15 | 0.56 | -0.54 | 0.41 | 0.37 | 0.35 | 0.57 | 0.88 | 1 |  |  |  |  |  |  |
| BIO14z | 0.04 | -0.27 | -0.09 | -0.04 | -0.09 | 0.07 | -0.11 | 0.15 | -0.08 | 0.03 | 0.04 | 0.53 | 0.15 | 1 |  |  |  |  |  |
| BIO15z**^†*^** | 0.42 | 0.14 | 0.38 | -0.31 | 0.34 | 0.33 | -0.22 | 0.21 | 0.36 | 0.36 | 0.39 | 0.11 | 0.49 | -0.65 | 1 |  |  |  |  |
| BIO16z**^†^** | 0.54 | -0.28 | 0.43 | -0.53 | 0.14 | 0.55 | -0.53 | 0.39 | 0.37 | 0.33 | 0.56 | 0.91 | 0.99 | 0.18 | 0.46 | 1 |  |  |  |
| BIO17z**^†*^** | 0.08 | -0.28 | -0.03 | -0.09 | -0.07 | 0.11 | -0.14 | 0.17 | -0.05 | 0.05 | 0.09 | 0.57 | 0.18 | 0.99 | -0.63 | 0.22 | 1 |  |  |
| BIO18z | 0.42 | -0.4 | 0.17 | -0.36 | 0.04 | 0.41 | -0.42 | 0.54 | 0.14 | 0.29 | 0.4 | 0.81 | 0.8 | 0.37 | 0.2 | 0.82 | 0.38 | 1 |  |
| BIO19z | 0.22 | -0.09 | 0.4 | -0.35 | -0.03 | 0.29 | -0.32 | -0.07 | 0.27 | 0.01 | 0.29 | 0.67 | 0.5 | 0.39 | -0.07 | 0.52 | 0.43 | 0.18 | 1 |

Variable definitions: BIO1 = annual mean temperature; BIO2 = mean diurnal range; BIO3 = isothermality; BIO4 = temperature seasonality; BIO5 = max temperature of warmest month; BIO6 = min temperature of coldest month; BIO7 = temperature annual range; BIO8 = mean temperature of wettest quarter; BIO9 = mean temperature of driest quarter; BIO10 = mean temperature of warmest quarter; BIO11 = mean temperature of coldest quarter; BIO12 = annual precipitation; BIO13 = precipitation of wettest month; BIO14 = precipitation of driest month; BIO15 = precipitation seasonality; BIO16 = precipitation of wettest quarter; BIO17 = precipitation of driest quarter; BIO18 = precipitation of warmest quarter; BIO19 = precipitation of coldest quarter.

**Supplementary Figures**

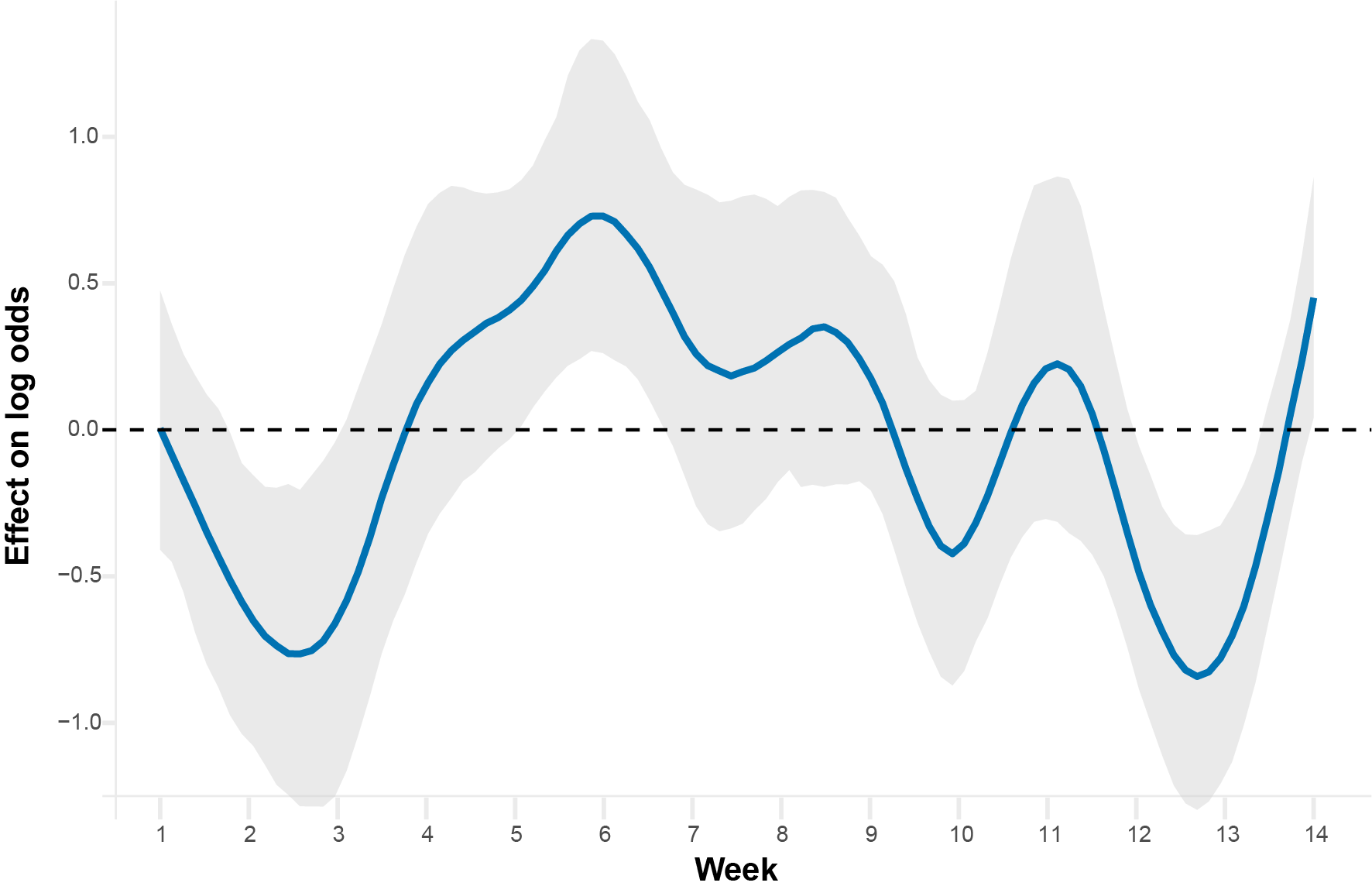

**Figure S1**. Partial effect of week on *w*Mel frequency estimated from a binomial generalized additive model (GAM), averaged across all 12 cages in the UPenn experimental orchard. The blue line shows the partial smooth on the logit scale and the shaded region represents the 95% confidence interval. The dashed horizontal line at zero indicates no partial effect of week on frequency. The x-axis spans 14 consecutive sampling weeks within a single season. The oscillating pattern suggests non-linear frequency changes across the duration of the experiment.

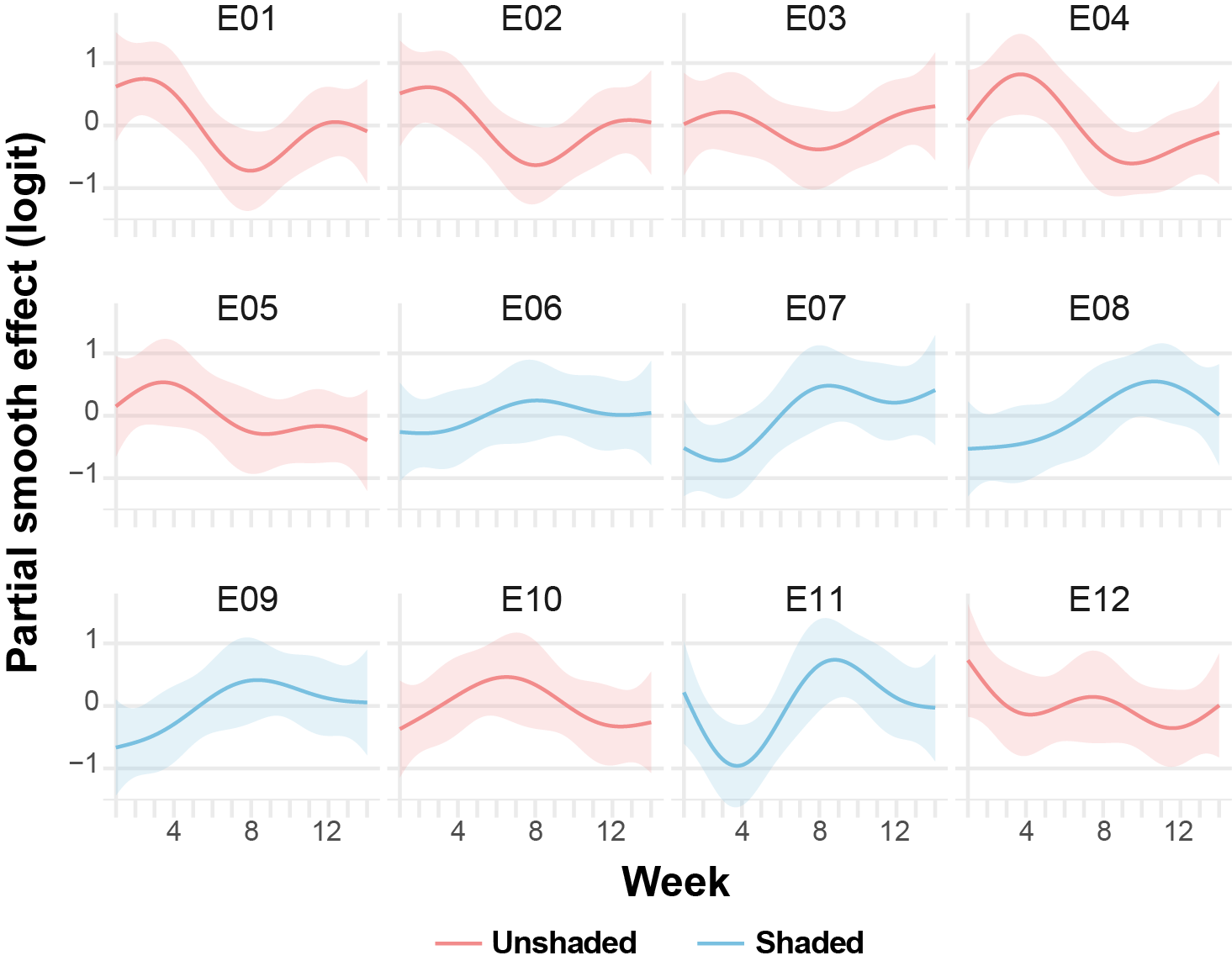

**Figure S2**. Partial effects of week on *w*Mel frequency for each of the 12 cages in the UPenn experimental orchard, estimated from a binomial generalized additive model (GAM). Each panel shows the cage-specific partial smooth on the logit scale across 14 consecutive sampling weeks within a single season, with shaded regions representing 95% confidence intervals. Curves are colored by *a priori* shade treatment: pink indicates unshaded cages and blue indicates shaded cages. The variation in smooth shape across cages reflects cage-level heterogeneity in within-season *w*Mel frequency dynamics, while the contrast between groups illustrates differences in temporal trajectories associated with shade classification.

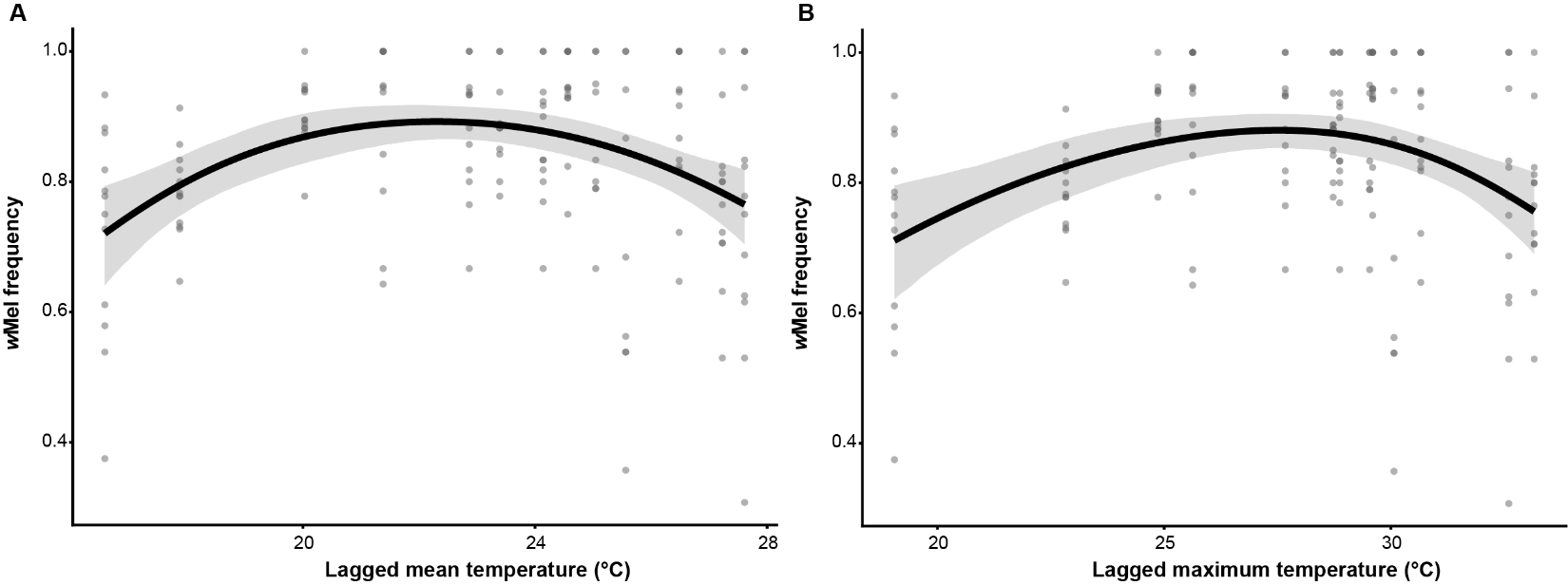

**Figure S3**. Relationship between temperature and *w*Mel frequency in the UPenn experimental orchard, estimated from binomial generalized additive models (GAMs). Each point represents an individual cage-week observation. **(A)** Effect of weekly mean temperature on *w*Mel frequency. **(B)** Effect of average weekly maximum temperature on *w*Mel frequency. In both panels, the curve shows the GAM-estimated smooth with 95% confidence interval (shaded region), and temperature values are lagged by one week to account for *Drosophila* developmental timelines. Both relationships show a unimodal response, with *w*Mel frequency peaking at intermediate temperatures and declining at temperature extremes.

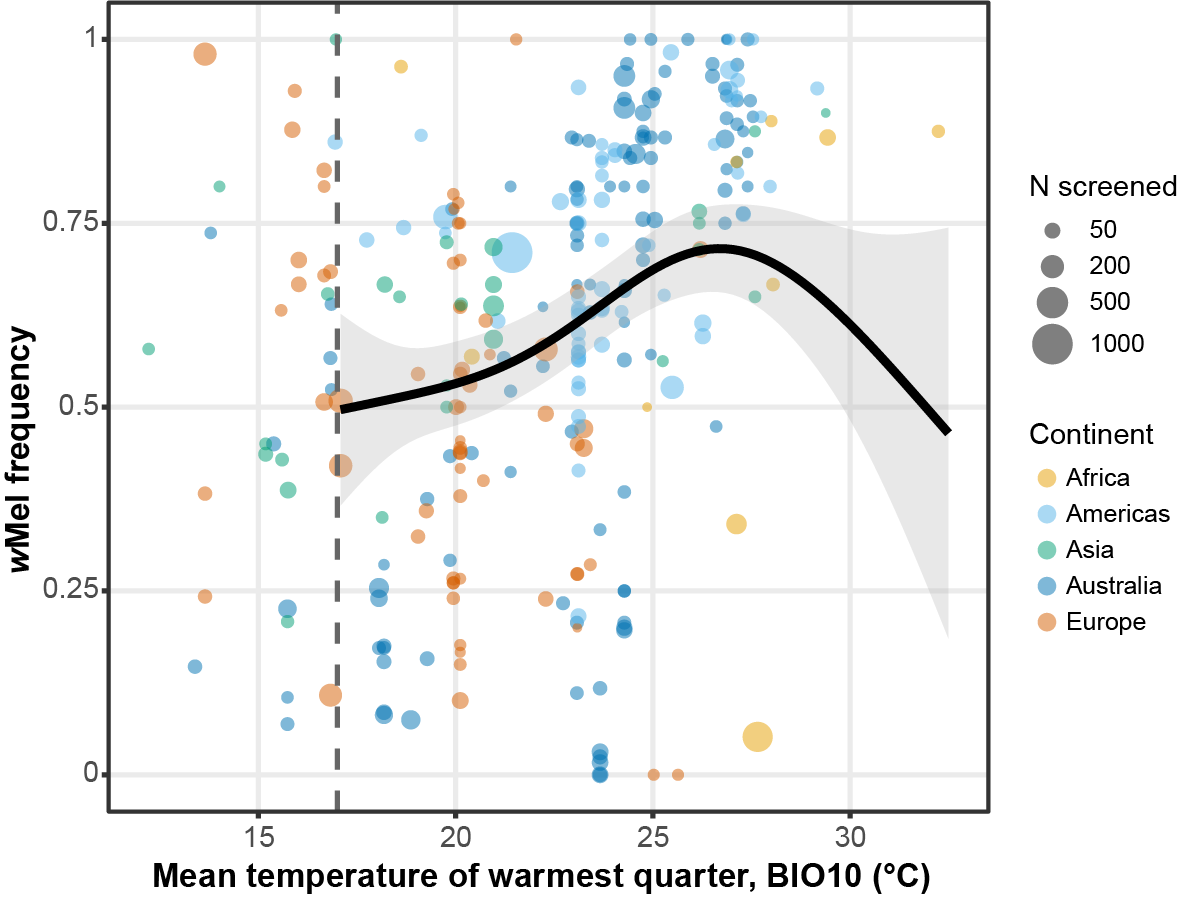

**Figure S4**. Raw *w*Mel frequencies as a function of mean temperature of the warmest quarter (BIO10) across global *D. melanogaster* locations. Each point represents a single spatiotemporal collection; circle diameter is proportional to the number of individuals screened for *w*Mel at that location, and colors indicate continent of origin. The vertical line at 18°C indicates the approximate temperature below which *D. melanogaster* populations are unlikely to persist year-round and may recolonize seasonally from lower latitudes or local refugia (Kriesner et al. 2016). BIO10 was not retained as a predictor in the bioclimatic model but captures the active *D. melanogaster* reproductive season more directly than annual mean temperature (BIO1); see main text for discussion.

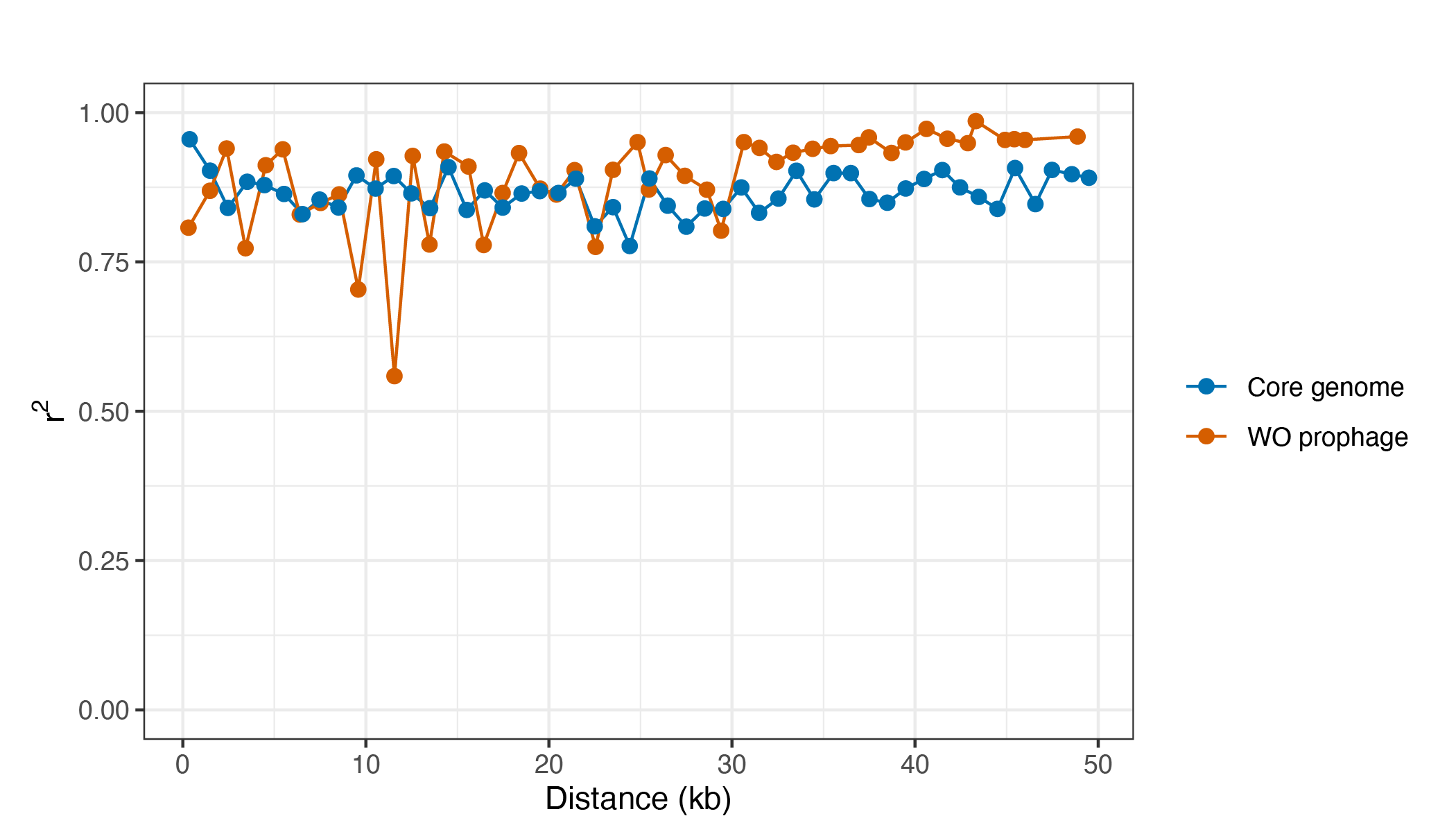

**Figure S5**. Linkage disequilibrium across the *w*Mel genome. Pairwise r² values between all biallelic SNPs plotted against physical distance (kb) for the core genome (blue) and WO prophage regions (orange). Linkage disequilibrium was near-complete (median r² = 0.94; median |*D*′| = 1.0) with little decay with physical distance (Spearman ρ between r² and distance = −0.042; |*D*′|: ρ = −0.029). Core and WO prophage regions plotted separately for presentation.

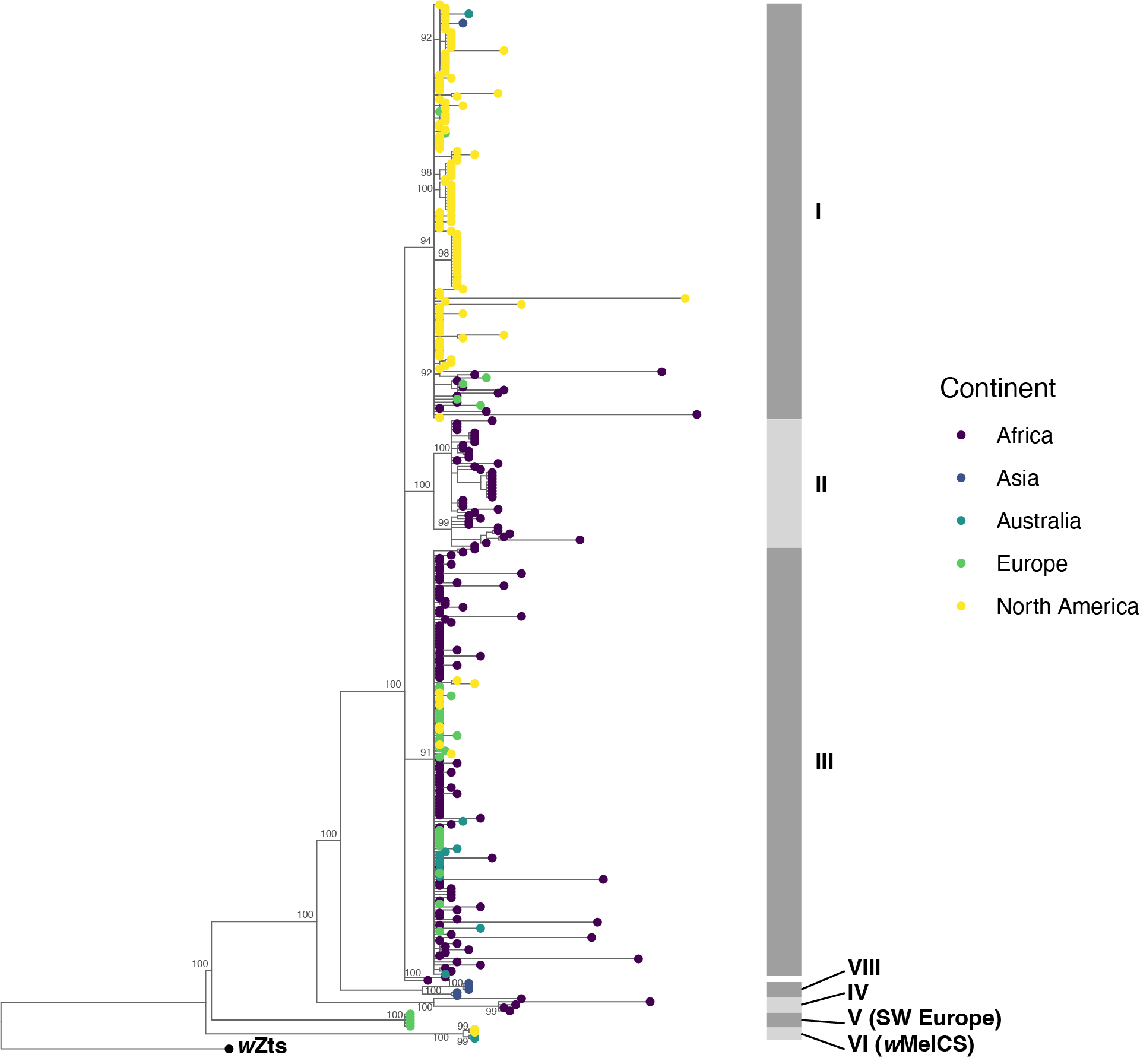

**Figure S6**. Maximum-likelihood phylogeny of all 339 individually sequenced *w*Mel genomes plus 14 reference sequences, based on 728 single-copy genes (736,557 bp) and rooted on *w*Zts from the sister species *Z. tsacasi* (Shropshire et al. 2026; 353 tips total). Tip circles are colored by continent of origin. Numbers at nodes indicate bootstrap support (%). Grey bars denote cytoplasmic clade designations following Richardson et al. (2012); alternating shading is used for visual clarity. Topology is identical to the pruned tree shown in Fig. 5A.

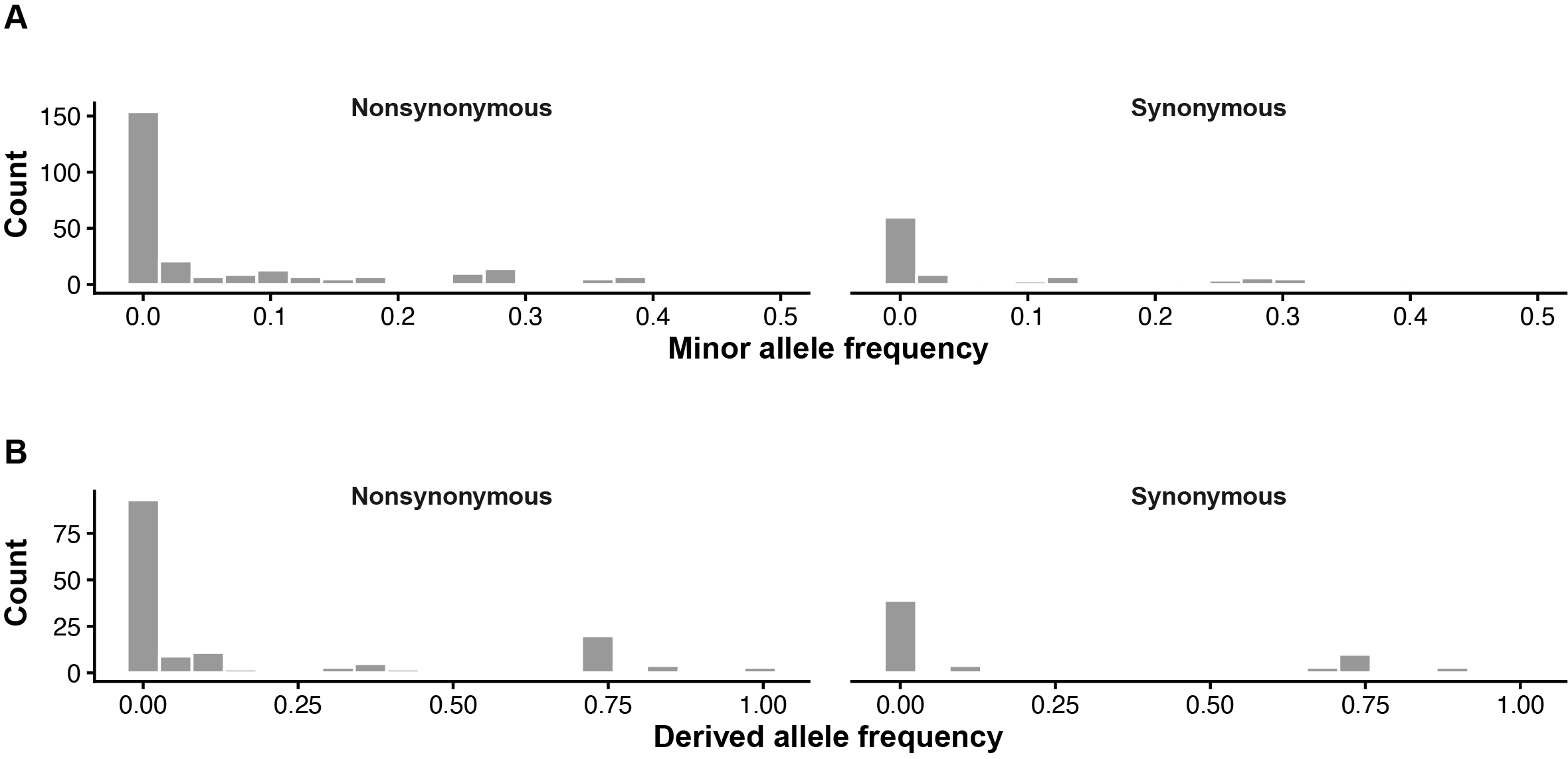

**Figure S7**. **Site frequency spectra of nonsynonymous and synonymous variants in the *w*Mel genome.** Each panel shows the distribution of allele frequencies for nonsynonymous (left) and synonymous (right) variants across all 593 biallelic SNPs, restricted to coding sites. **(A) Folded SFS.** Minor allele frequency (MAF) distributions for nonsynonymous (n = 271) and synonymous (n = 101) sites. Both distributions are strongly skewed toward rare alleles, and the two classes are indistinguishable (Kolmogorov-Smirnov test, D = 0.109, *P* = 0.67). **(B) Unfolded SFS, *w*Ri outgroup.** Derived allele frequency (DAF) distributions for 156 nonsynonymous and 65 synonymous coding sites polarized using Wolbachia strain *w*Ri as outgroup (221 of 372 coding sites polarizable). Both classes show a strong excess of low-frequency derived alleles (median DAF = 0.012 for both), and distributions did not differ significantly (KS test, D = 0.109, *P* = 0.65).

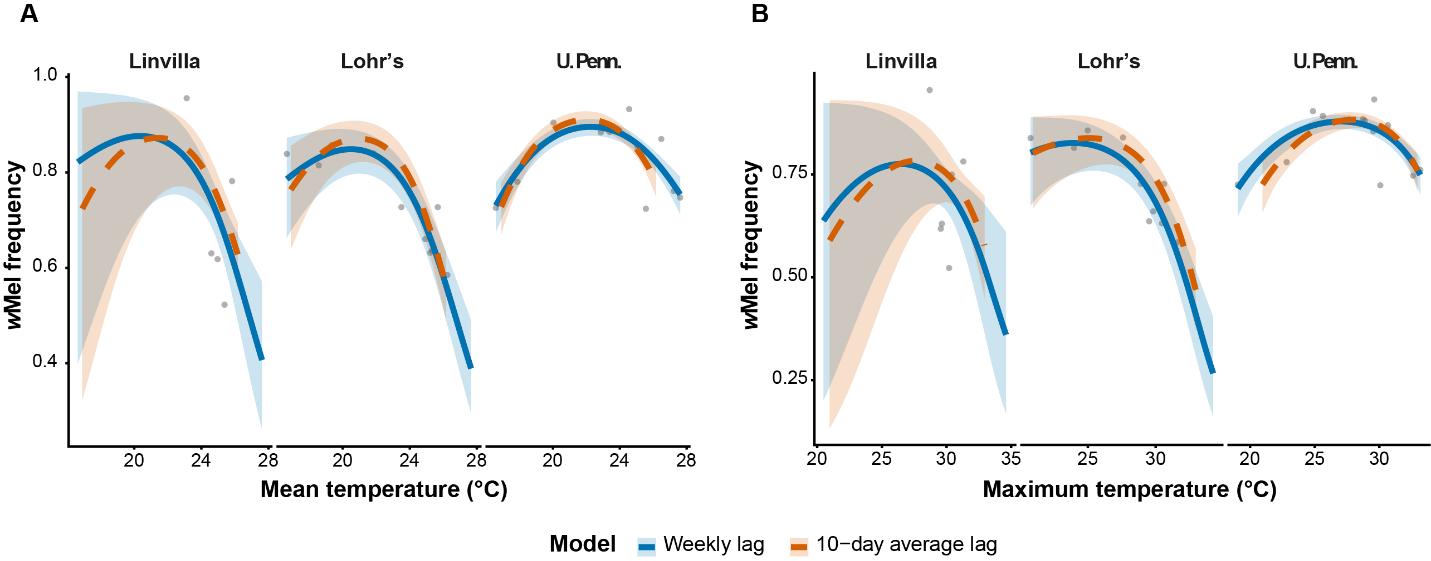

**Figure S8**. **Comparison of weekly lag and 10-day average lag temperature models predicting wMel frequency across three field locations.** Observed wMel frequency (grey points) and model-predicted wMel frequency (lines) as a function of (A) mean temperature and (B) maximum temperature across three locations (Linvilla, Lohr's, and U. Penn.). The blue solid line and shaded ribbon show predictions from the weekly lag model (lagged weekly mean and lagged weekly average maximum temperature, respectively) and the orange dashed line and shaded ribbon show predictions from the 10-day average lag model (mean temperature averaged over the 10 days prior to each observation). Shaded ribbons represent 95% confidence intervals. The two models produced highly correlated predicted wMel frequencies (mean temperature: Pearson's r = 0.91; maximum temperature: Pearson's r = 0.95).

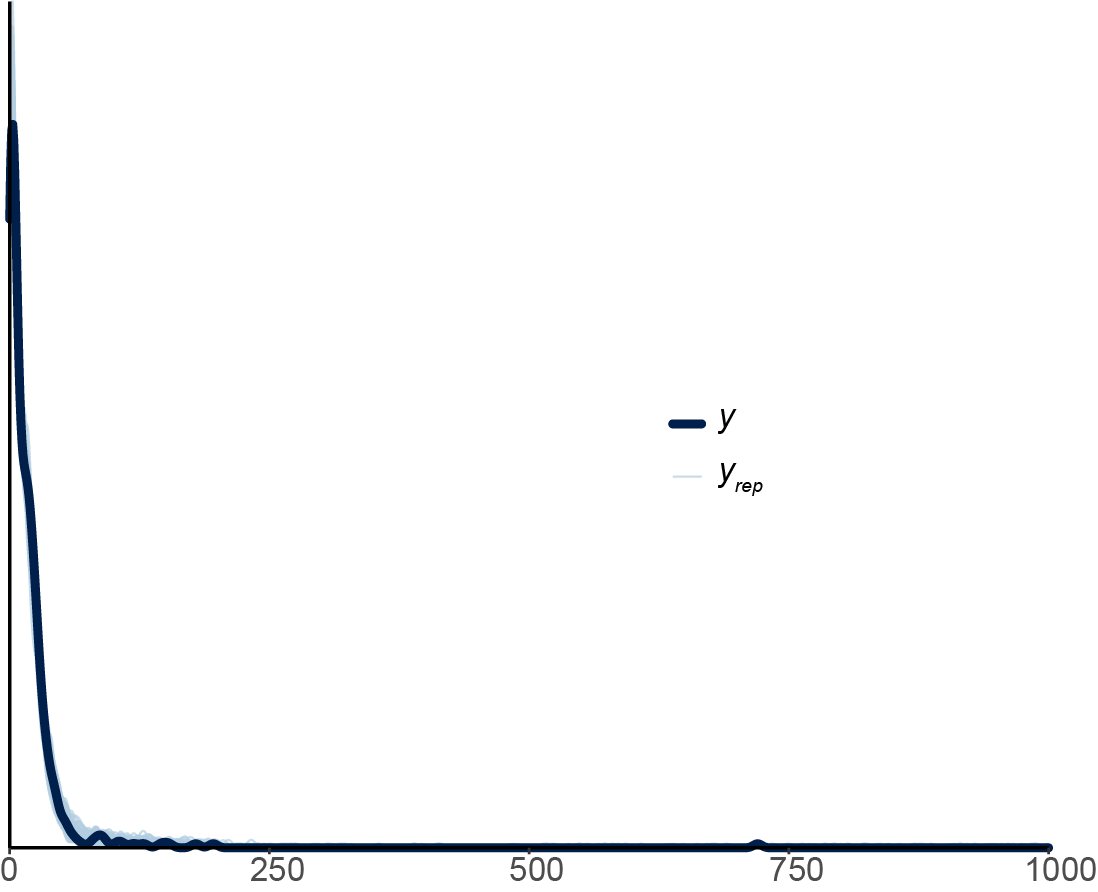

**Figure S9**. Posterior predictive check for the bioclimatic model with paired *w*Mel positive and total counts the dependent variable and residuals of scaled BIO6, scaled BIO8, scaled BIO15, and scaled BIO17 as global fixed effects, continent as a random intercept, and a continent-specific random slope for BIO8 estimated without intercept-slope correlation. 100 draws from the posterior predictive distribution are in blue. The observed data is represented in black. Overlap between observed and predicted distributions indicates adequate model fit.
